# Global, asynchronous partial sweeps at multiple insecticide resistance genes in *Aedes* mosquitoes

**DOI:** 10.1101/2024.04.09.588653

**Authors:** Thomas L Schmidt, Nancy M Endersby-Harshman, Anthony RJ van Rooyen, Michelle Katusele, Rebecca Vinit, Leanne J. Robinson, Moses Laman, Stephan Karl, Ary A Hoffmann

## Abstract

*Aedes aegypti* (yellow fever mosquito) and *Ae. albopictus* (Asian tiger mosquito) are globally invasive pests that confer the world’s dengue burden. Insecticide-based management has led to the evolution of insecticide resistance in both species, though the genetic architecture and geographical spread of resistance remains incompletely understood. This study investigates partial selective sweeps at resistance genes on two chromosomes and characterises their spread across populations. Sweeps at the volt-age-sensitive sodium channel gene (VSSC) on chromosome 3 correspond to one nucleotide substitution in *Ae. albopictus* and three substitutions in *Ae. aegypti*, including two at the same nucleotide position (F1534C) that have evolved and spread independently. In *Ae. aegypti*, we also identified partial sweeps at a second locus on chromosome 2. This locus contained 15 glutathione S-transferase (GST) epsilon class genes with significant copy number variation among populations and where three distinct genetic backgrounds have spread across the Indo-Pacific region, the Americas, and Australia. Local geographical patterns and linkage networks indicate VSSC and GST backgrounds probably spread at different times and interact locally with different genes to produce resistance phenotypes. These findings highlight the rapid spread of resistance genes globally and are evidence for the critical importance of GST genes in resistance evolution.

## Introduction

Insecticide resistance (hereafter: ‘resistance’) in mosquitoes and other insect pests remains one of the most pressing issues in global public health and food security, and the mechanisms underpinning its evolution continue to be a major focus of research ^1,2^. Genes associated with resistance include those conferring target-site resistance (e.g., mutations in the voltage-sensitive sodium channel (VSSC) gene ^3^) or metabolic resistance (e.g., glutathione S-transferase (GST) ^1^ and cytochrome P450 (CYP) genes ^4^). Genomic research on pests has helped identify how different resistance profiles are produced and maintained, such as by linkage, epistasis, and structural variation ^5–8^, and can indicate how specific types of resistance become geographically widespread via gene flow between populations ^9^ or species ^10^ or by multiple independent substitutions of the same allele ^11,12^. As resistance spreads it can produce striking patterns of geographical genetic structure in response to local selection pressures ^13^; this ‘local’ environment is determined by local insecticide usage, and thus geographically distant populations may have identical resistance alleles while proximate populations do not ^9,14^. Genetic structure at resistance genes can contrast sharply with genome-wide structure, which in invasive pests with large populations is generally established and reinforced by demographic processes during and after colonisation ^15^.

*Aedes aegypti* (yellow fever mosquito) and *Ae. albopictus* (Asian tiger mosquito) are highly invasive pests with a global dengue burden of ∼390 million infections per year^16^. Both species spread by human conveyance and have colonised much of the world’s tropics and subtropics and some temperate regions ^17,18^. In recent decades, incidental human movement has helped spread resistance alleles between populations of both species ^9,12,19,20^. As the global invasion of *Aedes aegypti* started in Africa and was mostly completed between the 16^th^ and 19^th^ centuries ^21^, the spread of resistance alleles in this species will mostly have involved gene flow between long-established populations. The invasion history of *Ae. albopictus* before the 20^th^ century is less well understood, though the species has since spread from its native range in Asia to colonise the Americas, the Pacific region, and Europe ^22^. *Aedes albopictus* may thus also be spreading resistance alleles as it invades new areas.

Evolutionary investigations of resistance in *Aedes* have predominantly focused on the VSSC gene, as resistant phenotypes can be traced to variation at this single gene. Several point mutations at this gene have been associated with resistance, but F1534C and V1016G are the two most well-characterised and widely distributed. F1534C has a global distribution in both species ^9,12^, while V1016G is common among *Ae. aegypti* in the Indo-Pacific region where it may be locally fixed or segregating alongside F1534C ^9^. V1016G has also recently been reported in *Ae. albopictus* ^23^. F1534C and V1016G confer strong resistance to Type I pyrethroid insecticides (e.g., permethrin), while V1016G also resists Type II pyrethroids (e.g., deltamethrin) ^3^. The advantages of VSSC mutations appear to be offset by physiological costs ^24^, which should favour the wild-type susceptible background when insecticide use is low. This trade-off has been observed in *Culex* mosquitoes, where local frequencies of resistant and wild-type backgrounds rise and fall in response to seasonal insecticide usage ^13^. In *Aedes* mosquitoes, this trade-off may also explain why wild-type mosquitoes can remain common in populations where VSSC resistance alleles are present ^9^ and why these alleles remain absent in some regions with low insecticide usage ^25^.

Genomic studies of *Ae. aegypti* have found that the F1534C mutation has likely evolved via two independent substitutions which have each spread globally ^12,19^. The V1016G allele appears to be from a single substitution and is usually observed with a third mutation, S989P ^9^. It is rarely observed on the same haplotype as the F1534C mutation ^9^. These alleles spread across populations by positive selection, which produces a selective sweep pattern in which individuals sharing that allele will also be more genetically homogeneous at sites near the “swept” locus than they are at other regions of the genome ^26^. A previous genomic study found selective sweeps through *Ae. aegypti* populations had structured genetic variation at sites more than 10 Mbp from the VSSC gene ^20^.

Genomic regions undergoing selective sweeps typically display three key evolutionary patterns: an increase in linkage disequilibrium, an increase in the proportion of alleles at low frequency, and an increase in population genetic differentiation ^27^. Within a single sample of individuals, these increases can be identified relative to other genomic regions. If sweeps are incomplete, or “partial”, they can be still more powerfully analysed using a second sample with a different swept background or no selection history at the locus of interest ^28^. If two different genetic backgrounds have undergone independent sweeps at the same locus, the two samples should be highly differentiated at the locus and this locus should show similarly high levels of linkage disequilibrium and rare allele proportions in each swept background. When one sample has experienced a sweep and one has not, these should be differentiated at the sweep locus and this locus should have higher levels of linkage disequilibrium and rare allele proportions in the sweep sample relative to the other sample. These patterns should be particularly stark if the sweep extends over populations strongly differentiated at other genomic regions.

This study analyses global patterns of genome-wide genetic structure in *Ae. aegypti* and *Ae. albopictus*, and detects SNPs that are structured in line with genotypes at VSSC point mutations rather than genome-wide structure. We use these SNPs to confirm two evolutionarily independent, partial sweeps of the F1534C allele and one of the V1016G allele in *Ae. aegypti*, and show for the first time a single sweep of the F1534C allele in *Ae. albopictus*, while greatly expanding current geographical knowledge of these sweeps and showing how these have spread across distant populations. However, our analysis also detected another genomic region in *Ae. aegypti* which has undergone three evolutionarily independent, partial selective sweeps, each segregating throughout multiple populations. Comparisons with other individuals from these populations showed the sweep-associated individuals had increased linkage disequilibrium, a greater proportion of low-frequency alleles, and stronger differentiation around the sweep locus, with similar patterns to those at the VSSC gene. This second sweep locus contained 15 GST epsilon class genes, with significant variation in GST gene copy number across countries. While GST epsilon genes have been linked to resistance to insecticides including DDT, organophosphates, and pyrethroids in *Aedes* and other mosquitoes ^29–31^, they have often been operationally treated as having partial contributions to metabolic resistance phenotypes alongside the many esterase and mono-oxygenase genes as well as other GSTs ^32^. The three evolutionarily independent, globally segregating backgrounds at the GST epsilon genes instead suggest these genes have a critical role in resistance to chemical control of *Ae. aegypti*.

## Results

### 2.1 Microhaplotype differentiation reveals global genetic structure and admixture patterns

Prior to investigating patterns of gene flow at specific genomic loci, we first investigated patterns of genetic structure and admixture across the genome. For this we used double digest restriction-site associated DNA sequencing (ddRADseq) data from 934 *Ae. aegypti* and *Ae. albopictus* individuals, including 358 sequenced for this study. This included *Ae. aegypti* from 19 countries (n = 444, Table S1) and *Ae. albopictus* from 17 countries (n = 490, Table S2).

As ddRAD data are sequenced as microhaplotypes, we investigated global genomic structure from patterns of haplotype coancestry, using fineRADstructure ^33^. This uses the sequence of all the SNPs from each RAD locus to find one or more closest relatives for each microhaplotype, producing coancestry matrices of individual haplotype similarity for *Ae. aegypti* (Fig 1a; 79,084 SNPs) and *Ae. albopictus* (Fig 2a; 96,269 SNPs). For both species, individuals formed clusters based on their population of origin and these populations tended to cluster by their geographical location. Notable exceptions were recent invasions of *Ae. albopictus* which clustered with their previously-identified source populations; these included Fiji (invaded from Southeast Asia ^34^), Mauritius (invaded from East Asia ^34^), and the Torres Strait Islands (invaded from Indonesia ^35^) (Fig 2a). Fig 2a also indicates that the recent *Ae. albopictus* invasion of Vanuatu originated in Papua New Guinea (PNG) or nearby.

**Fig 1.**
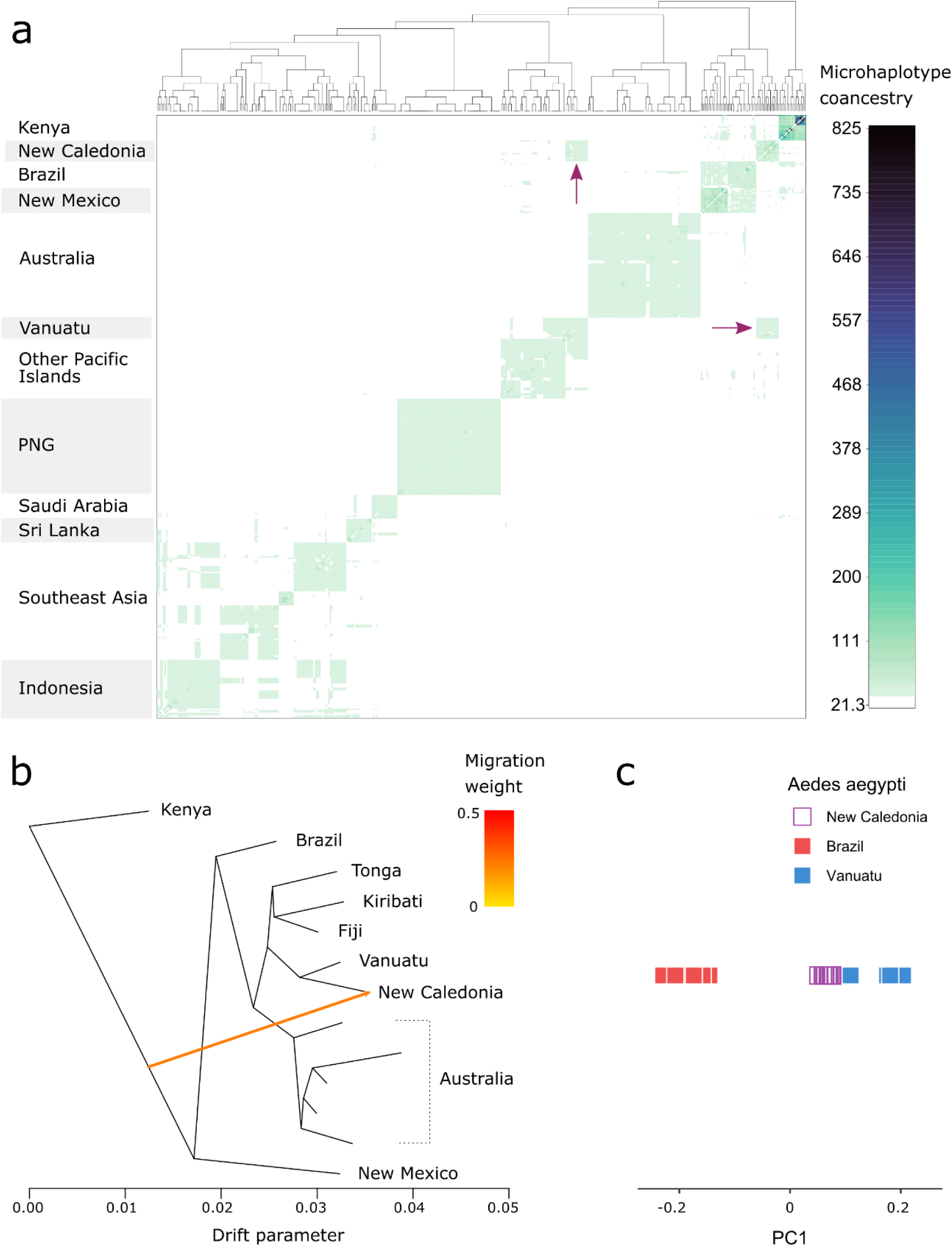
Genetic structure and admixture in *Ae. aegypti*. (a) Analysis of haplotype similarities among individuals in fineRADstructure ^33^. Darker colours indicate higher pairwise similarity of RADtag microhaplotypes. Arrows indicate putative admixture. (b) Maximum likelihood tree assessing admixture in TreeMix ^36^. TreeMix was run on windows of 500 SNPs using a subset of populations associated with the putative admixture identifed in (a). The coloured arrow indicates the migration edge. The single best tree of 100 is reported. (c) Principal components analysis of the putative admixed population and source populations. *Aedes aegypti* from New Caledonia are projected onto a single principal component constructed from variation between the two putative source populations. The position of the admixed individuals relative to the source populations reflects the admixture proportions ^39^.

**Fig 2.**
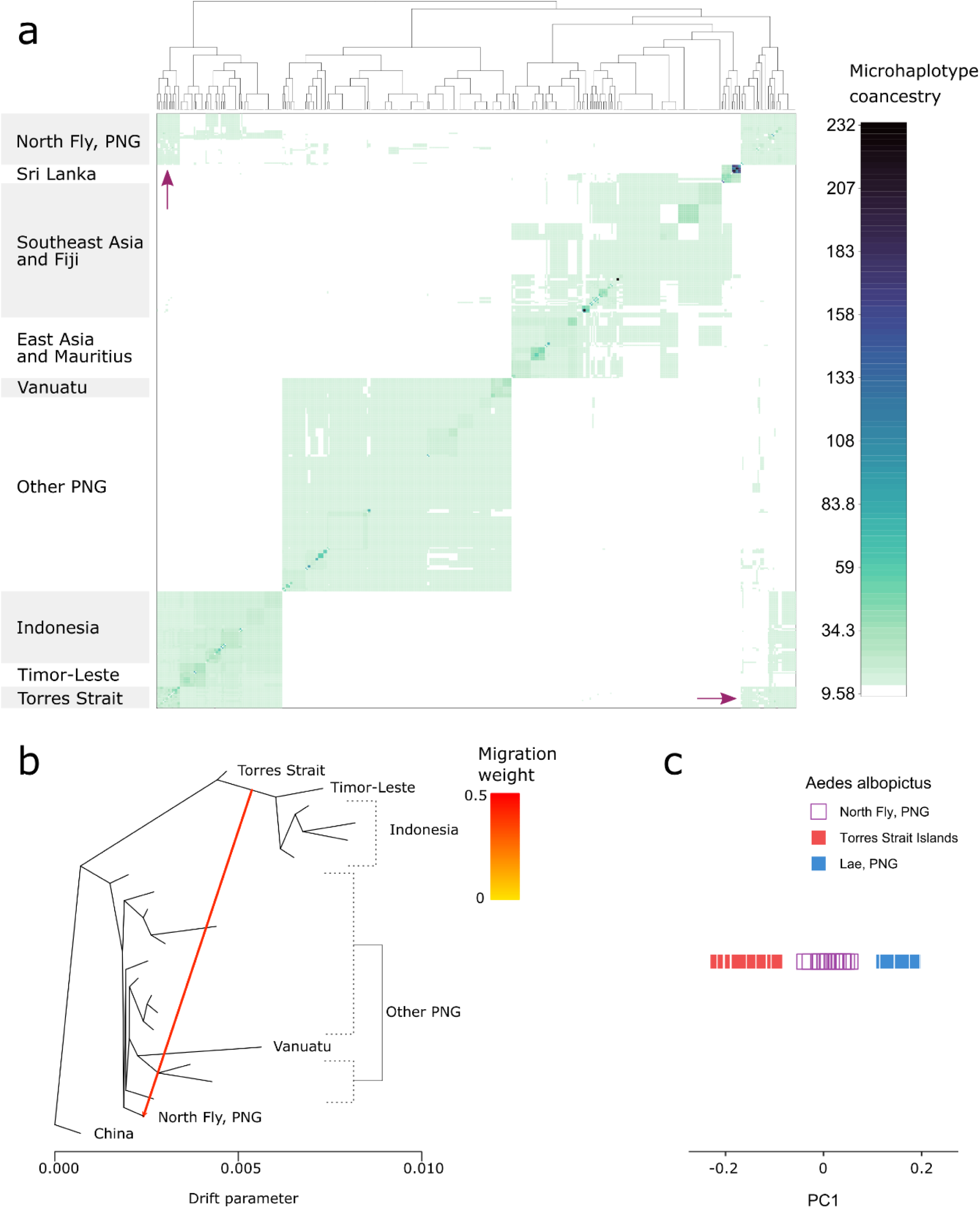
Genetic structure and admixture in *Ae. albopictus*. (a) Analysis of haplotype similarities among individuals in fineRADstructure ^33^. Darker colours indicate higher pairwise similarity of RADtag microhaplotypes. Arrows indicate putative admixture. (b) Maximum likelihood tree assessing admixture in TreeMix ^36^. TreeMix was run on windows of 500 SNPs using a subset of populations associated with the putative admixture identifed in (a). The coloured arrow indicates the migration edge. The single best tree of 100 is reported. (c) Principal components analysis of the putative admixed population and source populations. *Aedes albopictus* from North Fly, PNG, are projected onto a single principal component constructed from variation between the two putative source populations. The position of the admixed individuals relative to the source populations reflects the admixture proportions ^39^.

Putatively admixed populations were identified in *Ae. aegypti* from New Caledonia and *Ae. albopictus* from North Fly Region, PNG, which clustered apart from other populations in their regions (Fig 1a, 2a). We explored these relationships further using TreeMix ^36^ (Fig 1b, 2b), which builds maximum likelihood trees from population allele frequencies, then adds optional migration edges between populations. We ran TreeMix on subsets of populations selected from the fineRADstructure analysis, including putatively admixed populations, populations from potential source clades, and populations from clades intermediate to these. We assessed migration edge placement using 100 runs of TreeMix with either zero, one, or two migration edges added to the maximum likelihood trees. When TreeMix was run with one migration edge, these were placed in the same spots in all 100 TreeMix runs, and these trees had higher likelihood than the zero-migration trees in 98 (*Ae. aegypti*) and 99 (*Ae. albopictus*) of the 100 runs. When TreeMix was run with two migration edges, these were inconsistently placed across runs, so we used the tree with the highest likelihood and one migration edge. Here, *Ae. aegypti* from New Caledonia appeared to be an admixture of a population in the Americas or Africa (most likely the Americas given Fig 1a) and a population from the Pacific Islands (most likely Vanuatu) (Fig 1b), while *Ae. albopictus* from North Fly Region, PNG, was an admixture between other PNG populations and one or more populations from Indonesia, Timor-Leste, and the Torres Strait Islands (Fig 2b). We investigated these further using F_3_ statistics on all population triads. F_3_ statistics test whether one population has been produced from the admixture of two others, and are particularly powerful for detecting recent admixture events that involve equal proportions from each population ^37^, though may be underpowered for reduced representation data ^38^. F_3_ tests were statistically significant for admixture in North Fly Region, sourced from any of the other PNG populations and either the Torres Strait Islands or any Indonesian population (−3.95 × 10^-4^ < All F_3_ < -1.30 × 10^-3^, 0.031 > All Bonferroni P-values > 1 x 10^-23^). The most statistically probable pair of source populations were Bali, Indonesia and Madang, PNG. No F_3_ tests were significant for any *Ae. aegypti* populations, including comparisons involving New Caledonia. This result may reflect limitations of F_3_ tests in detecting older or asymmetrical admixture ^37^, which may be the case in New Caledonia, and may also reflect our limited sampling of American *Ae. aegypti* populations.

When admixed individuals are projected onto a principal component axis constructed from individuals from the two source populations, the position of the admixed individuals relative to the source population individuals reflects the admixture proportions ^39^. These PCAs indicated that New Caledonian *Ae. aegypti* were much more similar to Vanuatu than Brazil (Fig 1c), while *Ae. albopictus* from North Fly Region, PNG, had equivalent proportions of Torres Strait Island and PNG backgrounds (Fig 2c).

### 2.2 Three globally-distributed partial sweeps at the voltage-sensitive sodium channel (VSSC) in *Aedes aegypti*

To investigate selective sweeps, we aimed to identify genetic backgrounds at the VSSC gene that were common among geographically distinct populations of *Aedes* mosquitoes and that showed signs of positive selection. For this, we supplemented the genome-wide sequence data with endpoint genotyping assays that scored *Ae. aegypti* for three mutations at the VSSC gene (V1016G, F1534C, and S989P) and *Ae. albopictus* for two (V1016G and F1534C). In *Ae. aegypti,* all three mutations were detected, but as the S989P mutation was only found in the presence of V1016G we did not analyse it further. Genotypes at this site are recorded in Table S1. The V1016G and F1534C mutations were both geographically widespread and had similar global frequencies (0.333 and 0.327 respectively), though V1016G was not observed outside the Indo-Pacific region and neither mutation was found in any Australian population (Fig 3a). In *Ae. albopictus*, we did not detect any individuals with V1016G. After removing samples with uncertain VSSC genotypes, this left 395 *Ae. aegypti* and 490 *Ae. albopictus* for analysis. Frequencies of each allele in each population are listed in Tables S3 and S4.

**Fig 3.**
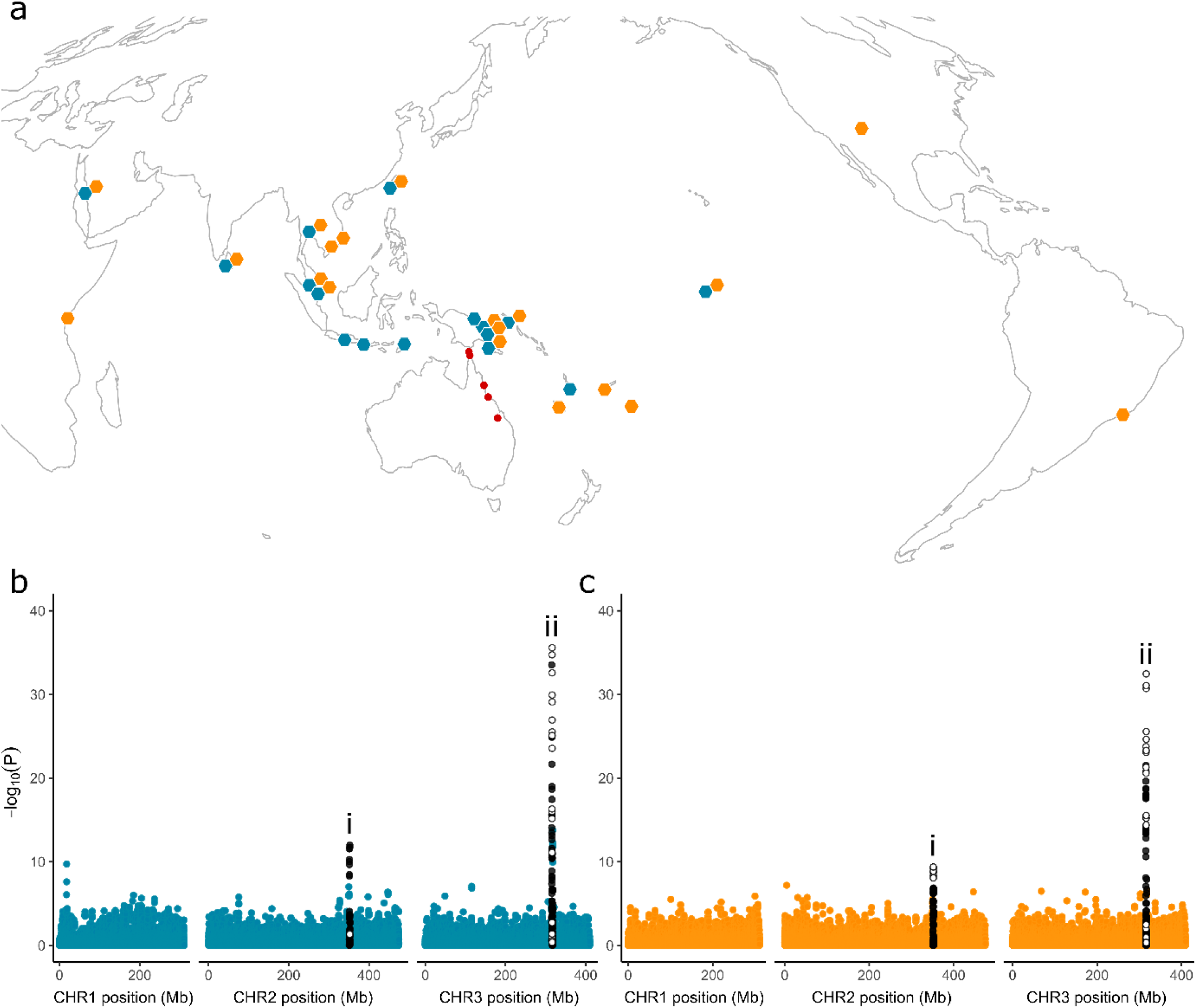
Geographical distributions of voltage-sensitive sodium channel (VSSC) mutations and their association with genome-wide SNPs in *Ae. aegypti*. (a) Coloured hexagons indicate populations containing the V1016G (teal) and F1534C (orange) mutations, with double hexagons indicating populations with both mutations. Red circles indicate populations where neither mutation was observed. Local frequencies of each mutation are listed in Table S3. (b, c) Latent factor mixed models for V1016G (b) and F1534C (c). Mixed models identify genome-wide SNPs that have genetic structure in line with VSSC genotype after controlling for genome-wide patterns assessed using sparse non-negative matrix factorisation with K = 18 (Fig S1). White circles indicate SNPs within the VSSC gene on chromosome 3 (‘ii’) or the region containing 15 glutathione S-transferase (GST) genes on chromosome 2 (‘i’), black circles indicate SNPs within 1 Mb of either region.

For *Ae. aegypti*, we first used latent factor mixed models to identify SNPs where genetic structure was segregating in line with genotypes at the V1016G and F1534C mutations rather than with genome-wide genetic structure. Genome-wide genetic structure was conditioned for in each mixed model via sparse non-negative matrix factorisation on the 51,115 genome-wide SNPs retained after filtering (Fig S1), setting K = 18. Outlier SNPs were identified as those with adjusted P-values below the inverse of the number of SNPs (1.956 × 10^-5^). We used these SNPs to identify genetic backgrounds shared across distinct populations, and assessed these for evidence of multiple independent substitutions of the same resistance mutation.

Mixed models identified 59 and 47 SNPs in and around the VSSC gene that were strongly structured in line with the V1016G (Fig 3b) and F1534C (Fig 3c) mutations. Strongly associated SNPs are listed in Table S5 and covered a region 2,555,236 bp overlapping the VSSC gene. Forty-two SNPs were common to both models (Fig 4a). Curiously, a second genomic region at ∼351 Mb on chromosome 2 also contained 13 (V1016G, Fig 3b) and 19 (F1534C, Fig 3c) SNPs strongly structured by VSSC genotype. This region contained 15 glutathione S-transferase (GST) genes and is analysed in section 2.3. Associations of SNPs with this second region appeared to be driven by the VSSC wild-type individuals from Australia, and rerunning the mixed models with these removed found no associations with the GST region, though the F1534C mutation was associated with a ‘Nach’ sodium channel protein (Fig S2).

**Fig 4.**
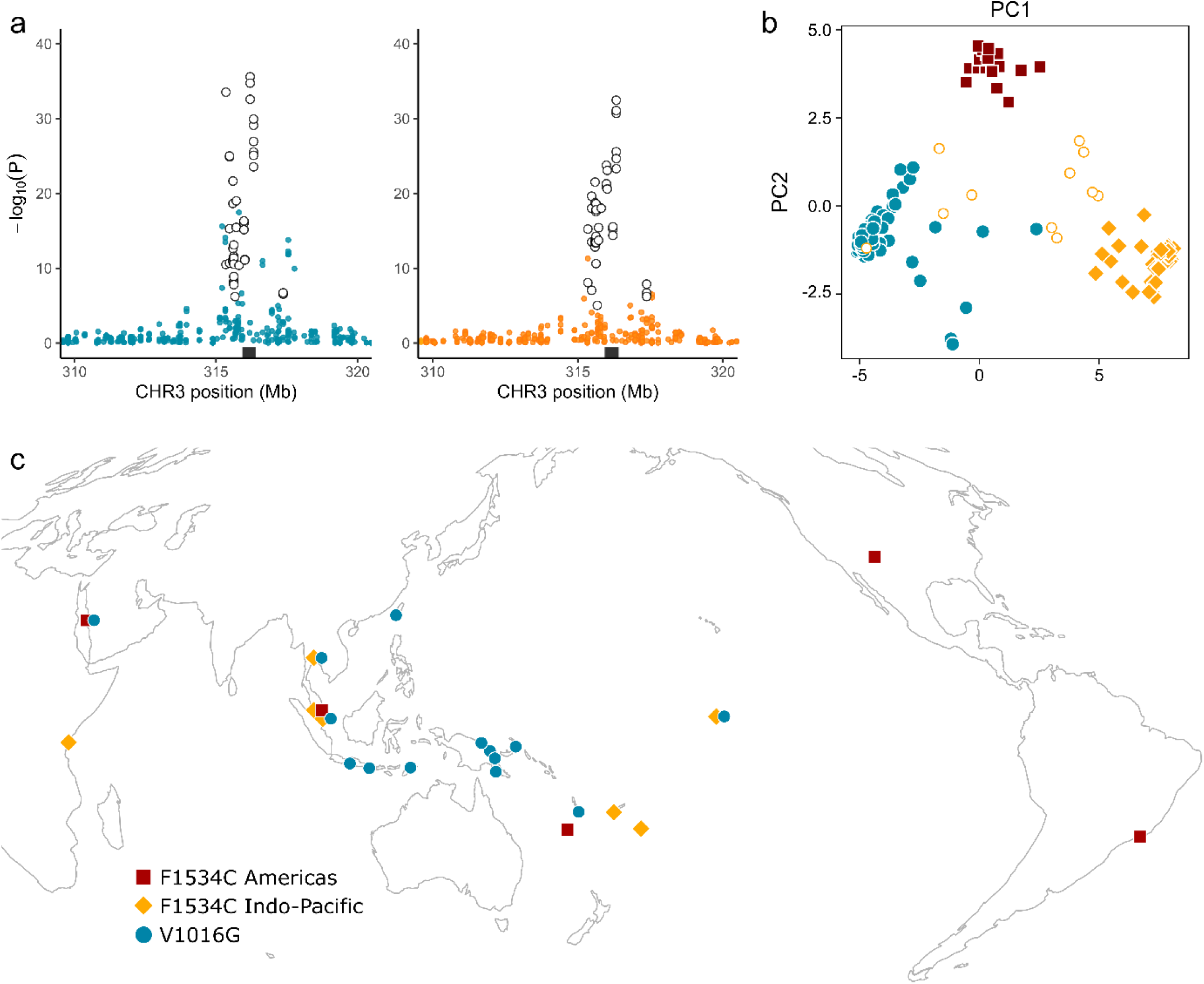
Identification of three sweep backgrounds at the voltage sensitive sodium channel (VSSC) gene in *Ae. aegypti*. (a) Latent factor mixed model results from Fig 3 in close-up. Black rectangles indicate the VSSC gene region. White filled circles indicate 42 SNPs identified by both models beyond the P < 1.956 × 10^-5^ significance cut-off. (b) PCA of all VSSC homozygotes, using the 42 outlier SNPs. Symbols of solid colour indicate homozygotes of the three swept backgrounds. White filled circles indicate F1534C homozygotes of uncertain background. (c) Locations containing at least one *Ae. aegypti* homozygote assigned to one of the three VSSC sweeps. Colours and shapes indicate the three swept backgrounds.

We used PCA of the 42 SNPs common to both models to infer evolutionary patterns (Fig 4b). V1016G homozygotes formed a single main cluster (n = 98), though some were spread toward the centre of the PCA likely as a result of missing data or recombination. F1534C homozygotes formed two main clusters. One of these included individuals from North and South America, New Caledonia, Saudi Arabia, and Malaysia (n = 21; Fig 4c; dark red) and the other included Kenya, Thailand, Malaysia, Fiji, Tonga, and Kiribati (n = 43; Fig 4c; orange).

We tested the hypothesis that the V1016G cluster and two F1534C clusters represented three independent sweeps spread across multiple populations via positive selection. To do this, we first estimated Tajima’s D (D), nucleotide diversity (π), and linkage disequilibrium (LD, taken as r^2^, the squared correlation coefficient between genotypes) between homozygotes assigned to each sweep (Fig 4c) and calculated the difference between these parameter estimates and those from all VSSC wild-type individuals (n = 113). In each case, the region around the VSSC gene showed unique patterns for ΔD (Fig 5a), Δπ (Fig 5b), and ΔLD (Fig 5c). For each putative sweep, ΔD and Δπ were lowest at the VSSC locus, while ΔLD was higher and with a broader peak than other regions on chromosome 3. The V1016G sweep had the weakest LD signal (Fig 5c), possibly due to the inclusion of recombined individuals (Fig 4b).

**Fig 5.**
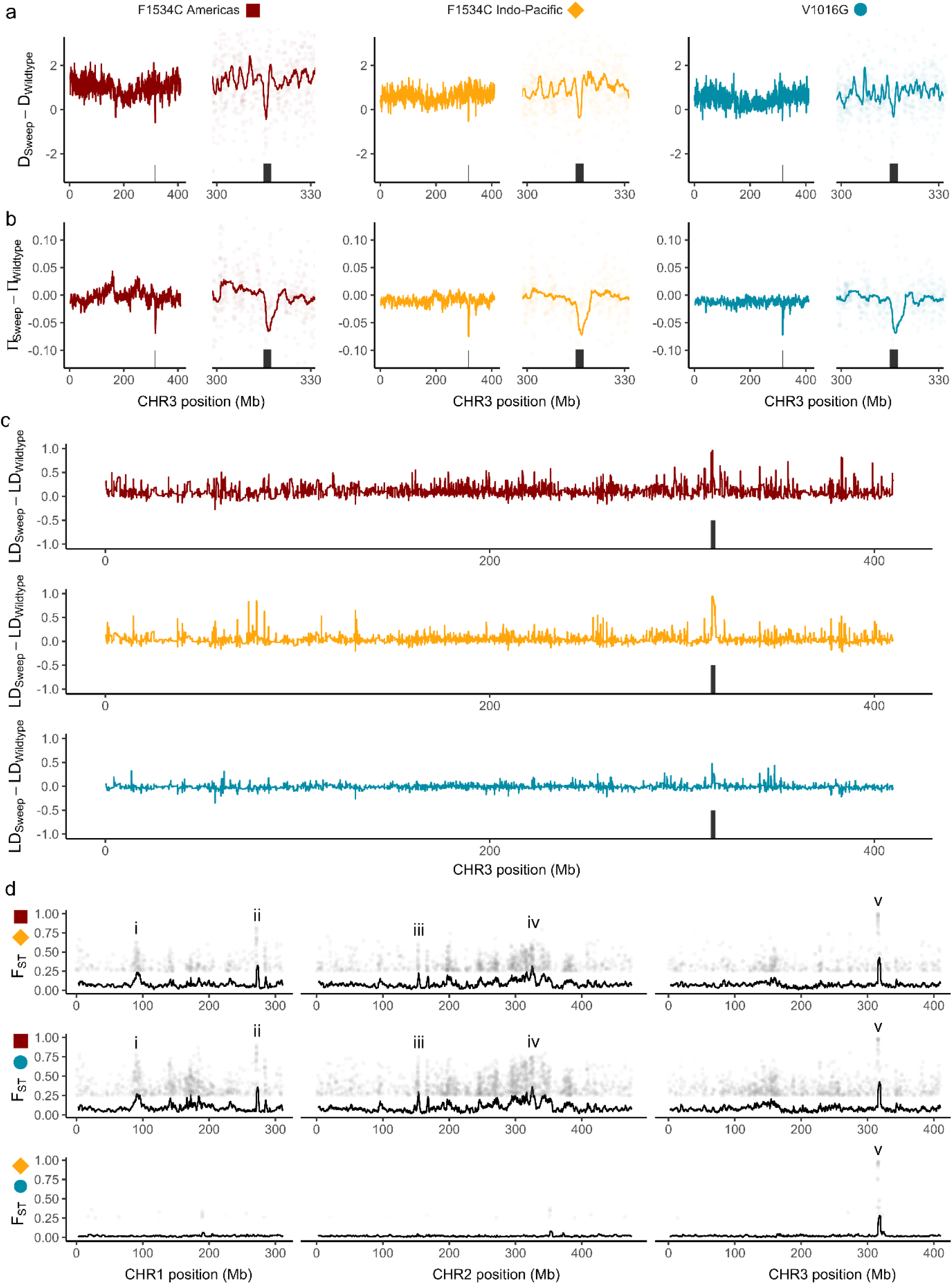
Genomic characterisation of three sweeps at the voltage sensitive sodium channel (VSSC) gene in *Ae. aegypti*. (a) Difference in Tajima’s D between sweep-associated individuals and wild-type individuals from all populations. Left hand plots show all of chromosome 3, right hand plots focus around the gene region. Black rectangles indicate the VSSC gene region ± 1 Mbp. Coloured lines show moving averages. (b) As above but showing differences in nucleotide diversity. (c) Moving average of the squared correlation coefficient between genotypes across chromosome 3. (d) Pairwise F_ST_ between individuals of each swept background. Colours and shapes indicate the two swept backgrounds being compared. Black lines show moving averages, plotted points show SNPs with F_ST_ > 0.25. Highly-divergent regions of interest are denoted ‘i’-‘v’ and described in the main text.

We calculated genome-wide pairwise F_ST_ between each pair of VSSC backgrounds to investigate these patterns further. Sharp F_ST_ peaks at the VSSC gene reflected many fixed differences between the three backgrounds (Fig 5d; ‘v’), suggesting these represent independent evolutionary events that have swept different alleles towards fixation. These analyses also indicated several other peaks across the genome (Fig 5d; ‘i-iv’) containing genes with functions relevant to insecticide resistance (complete list in Table S6). These include: (i) four ionotropic glutamate receptor genes with sodium transport function linked to insecticide resistance ^40^, and a probable glutathione peroxidase 2 gene with oxidative stress function linked to insecticide resistance ^41^; (ii) a sodium/potassium-transporting ATPase gene with sodium transport function, and a cytochrome P450 gene (CYP6BB2) with insecticide resistance function ^42^; (iii) a glutamate-gated chloride channel gene with insecticide resistance function ^43^; and (iv) an ankyrin gene with a function associated with the VSSC gene ^44,45^. The regions ‘i-iv’ with high F_ST_ were specific to comparisons between the F1534C Americas background (dark red) and the two Indo-Pacific backgrounds of F1534C (orange) and V1016G (teal).

### 2.3 Three partial sweeps and copy number variation at glutathione S-transferase epsilon genes (GSTs) in *Aedes aegypti*

The region of SNPs on chromosome 2 strongly associated with VSSC genotype covered a 2,761,683 bp region. This region contained 32 SNPs strongly associated (adjusted P-values < 1.956 × 10^-5^) with the V1016G and F1534C mutations (Fig 6a; 13 and 19 SNPs respectively). Genes within this region included 15 glutathione S-transferase (GST) epsilon-class genes with obvious links to resistance ^29–31^, as well as an ankyrin-3 gene with possible VSSC-associated function ^44,45^. Positions of the 32 SNPs are listed in Table S7.

**Fig 6.**
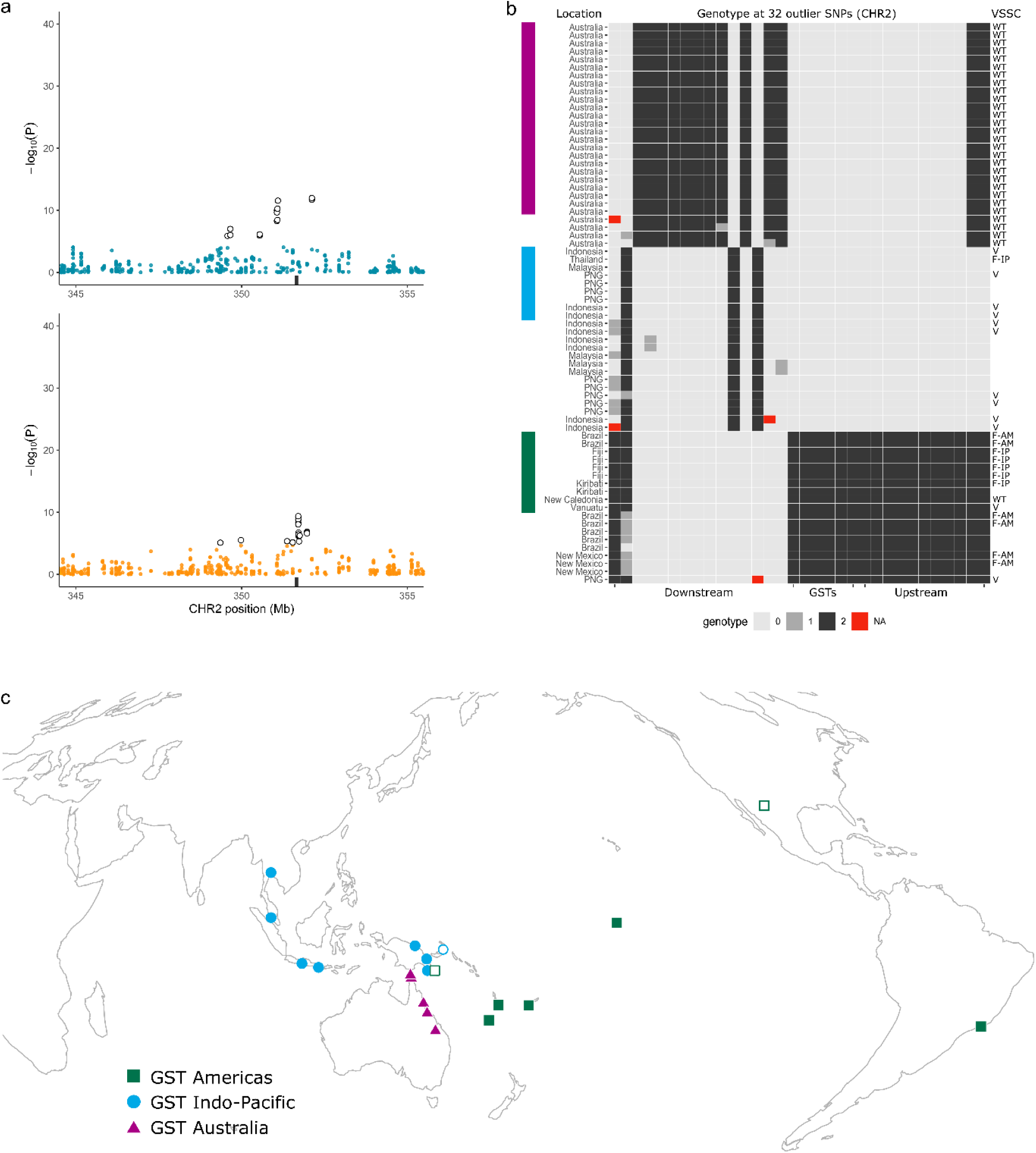
Identification of three sweep haplotypes at 15 glutathione S-transferase (GST) genes in *Ae. aegypti*. (a) Latent factor mixed model results from Fig 3 in close-up. Black rectangles indicate the region of 15 GST genes. White filled circles indicate 32 SNPs beyond the P < 1.956 × 10^-5^ significance cut-off. (b) Genotypes at the 32 outlier SNPs identified through latent factor mixed models. Each row represents one individual mosquito, with genotypes shaded by number of copies of the reference allele. Individuals indicated by the coloured rectangles have two copies of the same 32 SNP haplotype, other plotted individuals had up to one missing or heterozygous site. No other individuals are plotted. SNPs are split into those downstream, upstream, and within the region containing GST genes. Left hand side y-axis lists population of origin. Right hand side y-axis indicates VSSC genotype (WT = wild-type; V = V1016G background; F-A = F1534C Americas background; F-IP = F1534C Indo-Pacific background). (c) Locations of *Ae. aegypti* with two copies of a 32 SNP haplotype (coloured fill) or with one missing or heterozygous site (white fill). Colours indicate the three backgrounds following (b).

Visual analysis of variation at these SNPs identified three homozygous haplotypes in multiple individuals from different populations (Fig 6b). These were found in individuals from across all five Australian populations (GST Australia, purple, n = 24), from Thailand, Malaysia, Indonesia, and Papua New Guinea (PNG) (GST Indo-Pacific, blue, n = 9), and from Brazil, Kiribati, Fiji, New Caledonia, and Vanuatu (GST Americas, green, n = 10) (Fig 6c). VSSC genotypes of these 43 individuals were diverse, including homozygotes of V1016G (GST Indo-Pacific and GST Americas), F1534C IndoPacific (GST Indo-Pacific and GST Americas), F1534C Americas (GST Americas), and wild type (GST Australia and GST Americas) (Fig 6b). Although the three GST groups were identified by requiring all individuals to have homozygous genotypes at all 32 SNPs, relaxing this requirement to allow for up to one heterozygous or missing site added New Mexico and Port Moresby (PNG) to the GST Americas group. Port Moresby contained individuals from both the GST Americas and GST Indo-Pacific groups (Fig 6c).

We tested the hypothesis that the three GST haplotypes represented three independent sweeps spread across multiple populations via positive selection. Here we followed the same approach as for VSSC backgrounds (Fig 4), but, as the GST haplotypes were at lower frequencies than the VSSC mutations, there were enough wild types for us to restrict the ΔD, Δπ, and ΔLD comparisons to the same sets of populations as the sweeps: n = 47 (GST Australia), n = 89 (GST Indo-Pacific), and n = 62 (GST Americas). This should provide much more substantial evidence that the GST haplotypes were indicators of sweep backgrounds, and indicate whether the putative sweep backgrounds were segregating in populations (i.e. partial sweeps). In each case, the region around the GST genes showed unique patterns for ΔD (Fig 7a), Δπ (Fig 7b), and ΔLD (Fig 7c) that were similar to the VSSC gene. For each putative sweep, ΔD and Δπ were lowest at the GST locus, while ΔLD had a much broader peak than other regions on chromosome 2. Pairwise F_ST_ between each sweep background showed sharp peaks at the GST region in line with the differential fixation of alleles (Fig 7d; ‘ii’). A second peak was also apparent on chromosome 1 in the two comparisons with the GST Australia haplotype (Fig 7d; ‘i’). This contained a glutathione synthetase gene with antioxidant functions linked to insecticide resistance ^46^ (Table S8). Together these results suggest that the three haplotypes represent evolutionarily independent partial sweeps segregating in their populations.

**Fig 7.**
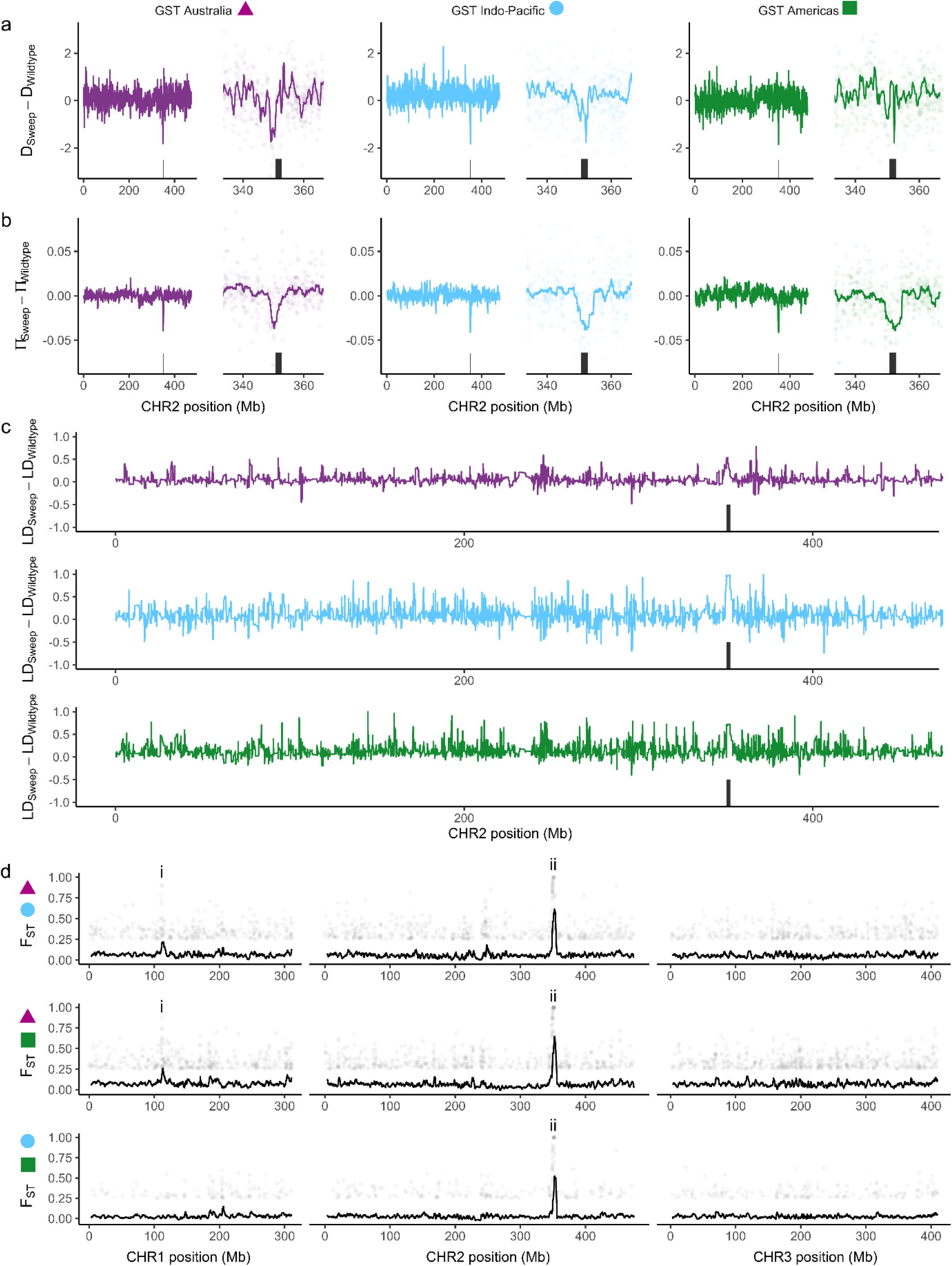
Genomic characterisation of three sweeps at 15 glutathione S-transferase (GST) genes in *Ae. aegypti*. (a) Difference in Tajima’s D between sweep-associated individuals and wild-type individuals from the same populations as the sweeps. Left hand plots show all of chromosome 2, right hand plots focus around the gene region. Black rectangles indicate the GST gene region ± 1 Mbp. Coloured lines show moving averages. (b) As above but showing differences in nucleotide diversity. (c) Moving average of the squared correlation coefficient between genotypes across chromosome 2. (d) Pairwise F_ST_ between individuals of each swept background. Colours and shapes indicate the two swept backgrounds being compared. Black lines show moving averages, plotted points show SNPs with F_ST_ > 0.25. The GST region is indicated with ‘ii’, the location of a glutathione synthetase gene is indicated with ‘i’.

Copy number variation among individuals was inferred through the ratio of read depths at sites within the coding regions of the 15 GST genes relative to the average depth at sites 10 Mb upstream and downstream from the GST region. Individuals with depth recorded at fewer than 500 sites within GST coding regions were omitted. For all individuals, downstream read depths were similar to upstream read depths (R^2^ = 0.98). Read depth ratios varied strongly by geographical region (ANOVA, F_9,263_ = 60.0, P = 1.1e^-58^) and were on average lower than upstream and downstream (x̂ = 0.86×) (Fig 8a). Ratios were much higher for individuals from North America (New Mexico, x̂ = 2.62×) but lower in the Middle East (Saudi Arabia, x̂ = 0.31×) and South Asia (Sri Lanka, x̂ = 0.30×); these two latter populations are genomically similar (Fig 1a). The higher read depths in New Mexico are unlikely to be from laboratory or sequencing errors, as these individuals were prepared and sequenced in the same DNA library as East Africa (Kenya). Read depth ratios showed no difference between individuals with GST sweep haplotypes and wild-type individuals from the same regions (Fig 8b). No signs of copy number variation were observed at the VSSC gene, though V1016G homozygotes had slightly higher relative read depths (Fig S3).

**Fig 8.**
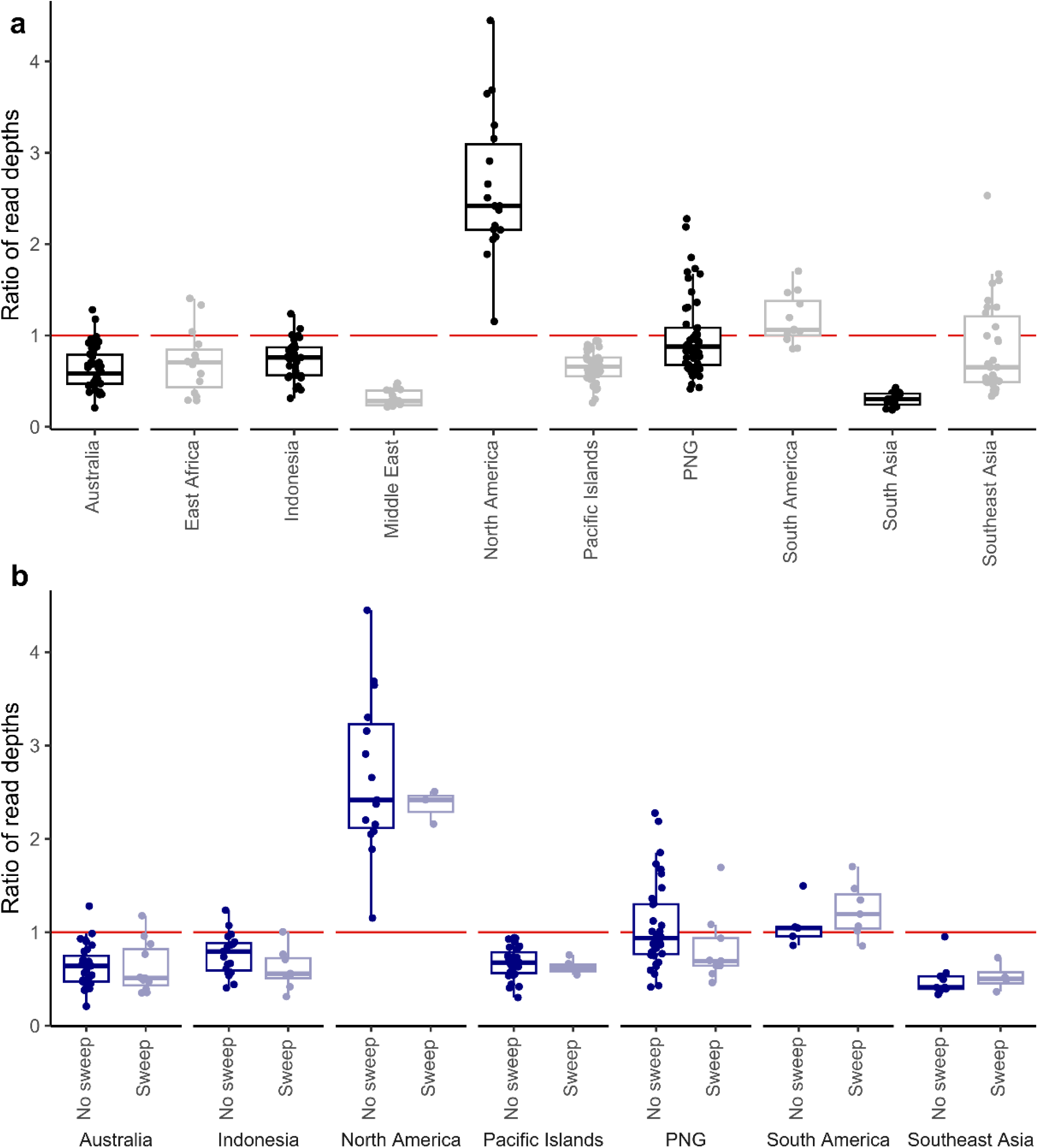
Copy number variation at 15 glutathione S-transferase (GST) genes in *Ae. aegypti*. Y-axis displays the ratio of read depths for each individual at coding regions in GST genes relative to sites <10 Mb upstream and downstream. Individuals with fewer than 500 sites scored within coding regions were omitted. Boxplot centre lines indicate medians. (a) All regions. (b) Regions containing GST sweeps, comparing individuals with sweep backgrounds to those with no sweep background, and omitting populations with no sweep backgrounds.

### 2.4 Linkage disequilibrium networks in *Aedes aegypti*

Linkage disequilibrium networks can provide useful insights into the genetic architecture of local adaptation ^47^. For each of the six sweep backgrounds in *Ae. aegypti* (three VSSC, three GST), we identified and investigated other regions of the genome that were in strong linkage (r^2^) with the sweep locus. As before, we analysed individuals in subsets corresponding to each sweep background and wild types. We considered only SNPs that were at least 50 Mb from the sweep locus and with r^2^ > 0.6 to at least one SNP within 1 Mb of the locus. SNPs were scored for each r^2^ > 0.6 interaction with a SNP near the locus, and these were plotted as histograms using 500 Kb bins (Fig 9).

**Fig 9.**
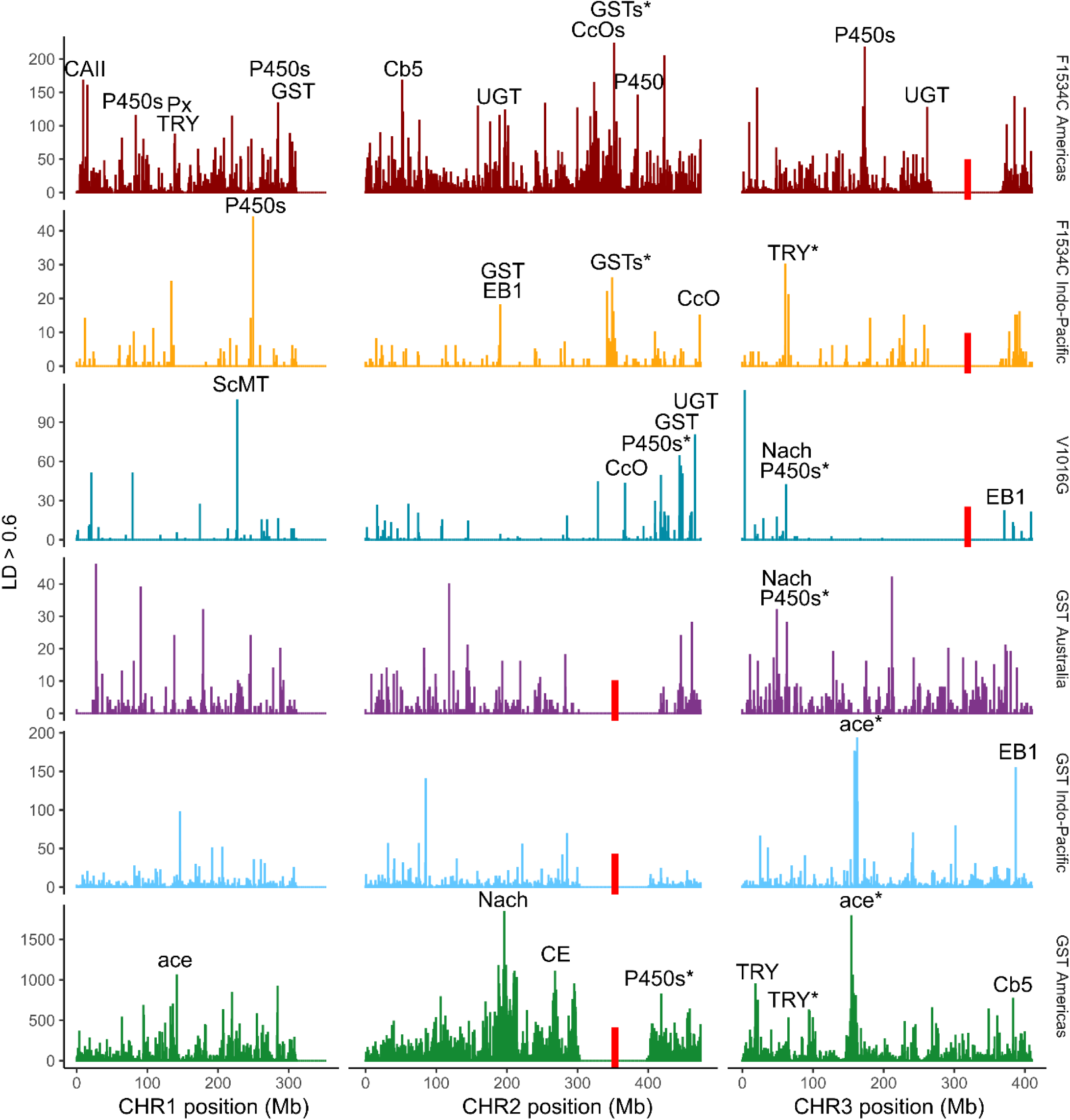
Linkage network analysis in *Ae. aegypti*. Rows indicate the three VSSC backgrounds (top) and the three GST backgrounds (bottom). Plots are histograms with 500 Kb bins, showing locations of SNPs with r^2^ > 0.6 to at least one SNP within 1 Mb of the sweep locus (red bars), and scoring SNPs for each r^2^ > 0.6 interaction with a SNP near the locus. SNPs within 50 Mb of the sweep locus were omitted. Labels indicate peaks containing gene(s) of known resistance association: P450 = cytochrome P450; GST = glutathione S-transferase; ace = acetylcholinesterase; UGT = UDP-glycosyltransferase; Nach = sodium channel protein Nach; Cb5 = cytochrome b5; CcO = cytochrome c oxidase; EB1 = esterase B1; CE = cholinesterase, CAII = carbonic anhydrase II; Px = peroxiredoxin-2; TRY = trypsin or anti-chymotrypsin; ScMT = sodium-coupled monocarboxylate transporter. Asterisks indicate peaks found in more than one background. Full list of genes in Table S9, raw linkage data in source data.

Peaks were observed for each of the six sweep backgrounds, and these overlapped genes with products linked to insecticide resistance, including cytochrome P450 ^4^, glutathione S-transferase ^1^, acetylcholinesterase ^6^, cytochrome b5 ^48^, cytochrome c oxidase ^49^, esterase B1 ^50^, Nach sodium channel protein ^3^, cholinesterase ^51^, UDP-glycosyltransferase ^52^, carbonic anhydrase II ^53^, trypsin and anti-chymotrypsin ^54^, peroxiredoxin ^55^, and sodium-coupled monocarboxylate transporter ^56^. Both F1534C backgrounds were strongly linked to the GST epsilon gene region, but no GST backgrounds were linked to the VSSC region. Several other genes were linked to multiple sweeps. These included a cluster of 19 cytochrome P450 genes on chromosome 2, a second cluster of 3 P450 genes and a Nach sodium channel protein gene on chromosome 3, an acetylcholinesterase gene on chromosome 3, and an anti-chymotrypsin gene on chromosome 3 (Fig 9). GST Australia had the fewest peaks associated with resistance genes. Wild-type individuals had far fewer strongly-linked SNPs and these formed smaller peaks at different locations to the sweep backgrounds (Figs S4-S5). We repeated the linkage network analyses on subsets of individuals from each background, either omitting countries or considering only specific countries (Figs S6-S10). These analyses indicated considerable heterogeneity in genetic architecture across countries and regions, where some genes were only associated with linkage peaks in specific regions and other genes were only associated with linkage peaks in the whole dataset. Genes identified through linkage network analysis are listed in Table S9.

### 2.5 A single partial sweep at the voltage-sensitive sodium channel (VSSC) in *Aedes albopictus*

We screened *Aedes albopictus* for the V1016G and F1534C mutations, finding no evidence of V1016G but six populations with F1534C (Fig 10a; Table S4). These were Singapore, Timor-Leste, Vanuatu, and three PNG populations. Frequencies of F1534C (0.033) were ten times lower than in *Ae. aegypti* (0.327), with only a single F1534C homozygote detected in PNG, and the North Fly PNG population had only a single heterozygous individual (n = 42 assayed; frequency = 0.012). We investigated signs of positively-selected variants across populations using a latent factor mixed model as in *Ae. aegypti*, though restricted our analyses to the six populations with VSSC mutations. Sparse non-negative matrix factorisation run on 45,809 SNPs conditioned for genome-wide genetic structure, setting K = 4 (Fig S11). A second latent factor mixed model run on all populations identified no regions of strong association (Fig S12).

**Fig 10.**
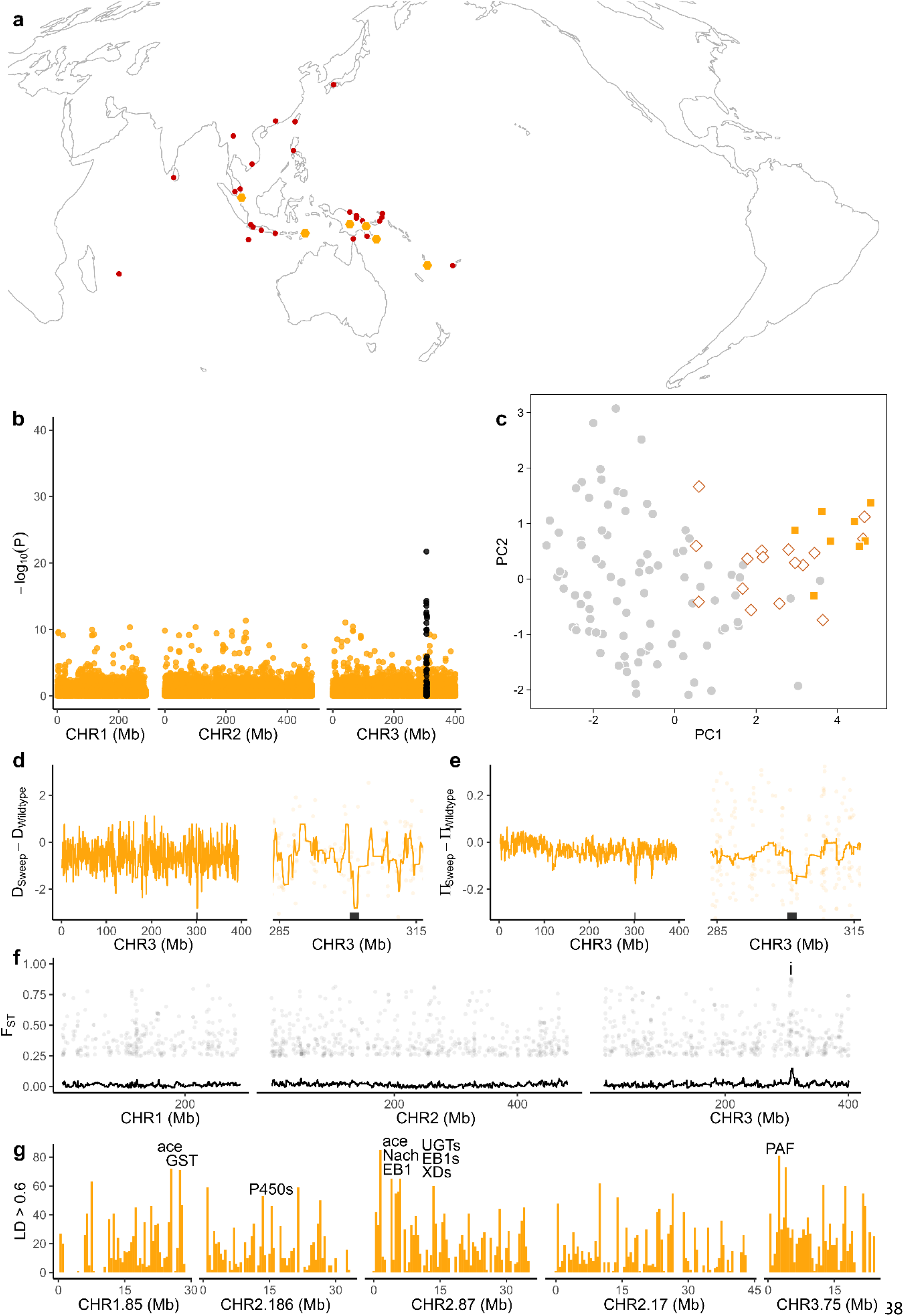
Genomic characterisation of the F1534C mutation at the voltage sensitive sodium channel (VSSC) gene in *Ae. albopictus*. (a) Orange hexagons indicate populations with F1534C, red circles indicate other populations. Local frequencies of each mutation are listed in Table S4. (b) Latent factor mixed model identifying genome-wide SNPs with genetic structure in line with VSSC genotype after controlling for genome-wide patterns assessed using sparse nonnegative matrix factorisation with K = 4 (Fig S11). Black circles indicate SNPs within 1 Mb of the VSSC gene on chromosome 3. Only the six populations with F1534C were included in the model. (c) PCA of 19 SNPs strongly associated with F1534C. Orange squares = F1534C homozygote, white filled lozenges = heterozygote, grey circles = wild type. (d) Difference in Tajima’s D between sweep-associated individuals and wild-type individuals from the same six populations as the sweeps. Left hand plot shows all of chromosome 3, right hand plot focuses around the gene region. Black rectangles indicate the VSSC gene region ± 1 Mbp. Coloured lines show moving averages. (e) As above but showing differences in nucleotide diversity. (f) Pairwise F_ST_ between individuals of the swept background and wild-type individuals from the same six populations. SNPs with F_ST_ > 0.25 are plotted. The VSSC region is seen as a peak on chromosome 3, indicated by ‘i’. (g) Linkage network analysis. Plots are histograms with 500 Kb bins, showing locations of SNPs with r^2^ > 0.6 to at least one SNP within 1 Mb of the sweep locus, and scoring SNPs for each r^2^ > 0.6 interaction with a SNP near the locus. SNPs within 50 Mb of the sweep locus were omitted. The five most strongly associated contigs are plotted. Labels indicate peaks containing genes of known resistance association: P450 = cytochrome P450; GST = glutathione S-transferase; ace = acetylcholinesterase; EB1 = esterase B1; UGT = UDP-glycosyltransferase; Nach = sodium channel protein Nach; XD = xanthine dehydrogenase; PAF = phenoloxidaseactivating factor 2. Full list of genes in Table S10, raw linkage data in source data.

The model identified 19 SNPs around the VSSC gene that were strongly associated with the F1534C mutation (adjusted P-value < 2.183 × 10^-5^) (Fig 10b). A PCA of these SNPs indicated clustering among individuals with the F1534C mutation, and homozygotes in particular, suggesting a shared evolutionary history and a single substitution event underlying the F1534C mutation in these samples (Fig 10c). Differences in Tajima’s D (Fig 10d) and π (Fig 10e) between F1534C homozygotes and wild-type individuals from the same populations showed similar signs of positive selection as in *Ae. aegypti* (Fig 5). Pairwise F_ST_ between F1534C homozygotes and wild-type individuals likewise indicated a peak at the VSSC region (Fig 10f).

Linkage network analysis was conducted with identical protocols to *Ae. aegypti* (section 2.4), though as the *Ae. albopictus* genome assembly is far less contiguous we focused on the five contigs with the largest number of r^2^ > 0.6 interactions. The five contigs all had peaks of strongly linked SNPs (Fig 10g), though with considerably more noise possibly due to small sample size (n = 8). Some peaks overlapped genes with products linked to insecticide resistance, including cytochrome P450 ^4^, glutathione Stransferase ^1^, acetylcholinesterase ^6^, UDP-glycosyltransferase ^52^, esterase B1 ^50^, Nach sodium channel protein ^3^, xanthine dehydrogenase ^57^, and phenoloxidase-activating factor ^58^. Genes identified through linkage network analysis are listed in Table S10.

Several genes identified by linkage network analysis were common to both *Ae. aegypti* and *Ae. albopictus*. This included a cluster of cytochrome P450 genes of types including 6a8, 6a14, 6a20, 6d3, and 6d4. These genes are found on chromosome 2 in both species, as a cluster of 19 genes in *Ae. aegypti* and 16 genes in *Ae. albopictus*. This cluster was linked to two backgrounds in *Ae. aegypti* (V1016G and GST Americas) and the F1534C background in *Ae. albopictus* (Figs 9, 10g). Both species also shared links to UDP-glucuronosyltransferase 2C1 genes found on chromosome 3 in *Ae. aegypti* and chromosome 2 in *Ae. albopictus*. These were linked to the F1534C Americas background in *Ae. aegypti*.

## Discussion

The spread of insecticide resistance in pests is a pressing global problem. Recent work in *Aedes* has shown how resistance mutations at the VSSC gene can reach global distributions via gene flow following multiple independent substitutions ^9,12,19^. Less is known about how metabolic resistance originates and spreads, though it is typically assumed to have more complex genetic architectures than target-site resistance ^32^. This study identified three distinct genetic backgrounds at GST epsilon genes that have spread by positive selection across *Ae. aegypti* populations. The global scale and population genetic patterns of these three genetic backgrounds are similar to the three VSSC backgrounds we also describe. While many different genes can contribute to metabolic resistance phenotypes, the observation that three evolutionarily independent backgrounds have undergone global partial sweeps at the same GST epsilon gene cluster strongly suggests this cluster has a major role in insecticide resistance. The global spread of GST backgrounds contrasts with those observed in anopheline mosquitoes where barriers to gene flow appear to have restricted the distribution of resistant alleles ^59^.

Although the GST and VSSC sweeps had similar broad geographical patterns, local patterns indicate these backgrounds spread mostly asynchronously. This is most evident in the Pacific Islands, where Vanuatu, Fiji, Kiribati, and New Caledonia all shared the same GST Americas haplotype, but each had only a single VSSC background of either V1016G, F1534C Indo-Pacific, or F1534C Americas (Fig 6). Vanuatu and Fiji were also at fixation and near fixation for these VSSC backgrounds (Table S3). However, this does not rule out the simultaneous spread of GST and VSSC backgrounds at some locations. Linkage network patterns add additional insight here, as both F1534C backgrounds were strongly linked to the GST epsilon region, but no GST backgrounds were linked to the VSSC gene (Fig 9). Additional temporal samples from key populations could help disentangle the shared evolutionary history of these genes.

The possible admixture patterns in New Caledonia provide additional insight into these invasion histories (Fig 1). If these patterns reflect gene flow from an American background into an established Pacific Island background, this could mark the point at which the GST Americas haplotype was brought into the Pacific Islands. As F1534C is at a moderate frequency in New Caledonia (p = 0.21), it is conceivable that the GST Americas haplotype spread from New Caledonia into Fiji, Kiribati, and Vanuatu by VSSC wild-type mosquitoes, though whether this occurred before or after the local introduction of VSSC mutations is unclear. New Caledonia was a hub of international shipping from the late 19^th^ century to WWII ^60^; given that *Ae. aegypti* rapidly reaches high local densities after colonisation, a large number of mosquitoes must have been introduced from the American source to produce genome-wide admixture patterns as strong as those observed ^15^. If admixture occurred before the 1960s, this likely predates the introduction of the GST Americas haplotype, as insecticidal control efforts only began in New Caledonia in the 1960s ^60^. There is also some confusion around the source and timing of the *Ae. aegypti* invasion into the Pacific Islands. While most evidence points to an invasion from the Mediterranean at the end of the 19^th^ century ^21^, an earlier invasion of the Pacific Islands from the Americas is plausible, given historical records of arboviral disease outbreaks in Southeast Asia ^61^ and early outbreaks of dengue in New Caledonia specifically ^60^. The PCA results and non-significant F_3_ tests suggest that if this hypothesis is true it may be too old or has experienced too much local gene flow to be detected ^37^.

Sweeps at the VSSC gene align with previous evidence that the F1534C allele is derived from two independent substitutions ^12,19^ and the fact that F1534C and V1016G have swept through the Indo-Pacific ^9^. Here, our extensive sampling provides a detailed indication of the extent to which each VSSC background has spread, most notably the F1534C Americas background, which was found not only in the Americas but at locations in the Pacific Islands, Southeast Asia, and the Middle East (Fig 4c). A recent study identified another VSSC mutation, L982W, that produced strong resistance phenotypes when combined with F1534C but has not been recorded outside of Vietnam and Cambodia ^62^. Here, F1534C was observed in *Ae. aegypti* samples from Ho Chi Minh City and Nha Trang, but only as a heterozygote (Table S1). If the L982W mutation is presently spreading through any other Indo-Pacific populations, it may be detectable as a third F1534C background, particularly if this background spreads through populations that are already at or near fixation for VSSC mutations. This assumes the new haplotype confers an additional selective advantage in wild populations.

Compared with *Ae. aegypti*, *Ae. albopictus* had far fewer populations with VSSC resistance alleles (Fig 10a, c.f. 4c), and when present these alleles were at lower frequencies (Tables S3,S4). Despite the comparative rarity of these alleles, a single F1534C allele copy was found in the most geographically remote population, in North Fly Region, PNG. F1534C was found at low frequency in three PNG populations >600 km apart and may be present in other unsampled locations such as in the South Fly Region, which shares a genetic background with Torres Strait Islands *Ae. albopictus* and is likely the source of the admixture into the North Fly Region ^35^. The spread of this allele across six populations points to a selective advantage, and low frequencies of this allele compared with those in *Ae. aegypti* might be due to limited insecticide exposure in sylvan habitats where *Ae. albopictus* is more common than *Ae. aegypti*^63^. The current geographical distribution of F1534C and its apparent selective advantage suggest future insecticide usage against *Ae. albopictus* may lead to F1534C rapidly increasing in frequency.

There was substantial variation in GST epsilon class read depth among regions, with North America ∼8.5 times higher than the Middle East and South Asia (Fig 8). Given the GST Americas haplotype was segregating in New Mexico, though only found as a heterozygote, this copy number increase likely occurred after the initial sweep. This inference aligns with the observation that none of the GST sweeps were linked to gene duplications (Fig 8b), and together these point to different mutational mechanisms underpinning resistance, such as point mutations; a single mutation in a GST epsilon gene was linked to metabolic resistance to DDT in Beninese *An. funestus* ^30^. The New Mexico population in this study is part of an *Ae. aegypti* genetic cluster extending from Southern USA through to Central America and the Caribbean ^64^, and it would be valuable to explore copy number variation and associated resistance further in this region. Positive selection was recently detected at this GST gene cluster in Brazil and Colombia ^65^.

The GST Australia sweep likewise demands further investigation. Usage of insecticides against mosquitoes in Australia has been low since DDT was banned in 1987^66^. The GST Australia sweep may represent a second selective sweep at this locus, with the region reverting to a wild-type background after an initial resistant sweep, as observed in *Culex* mosquitoes ^13^. If so, this would require a considerable fitness cost of the resistant allele like that reported for the F1534C mutation in *Ae. aegypti* ^24^. The partial nature of the GST sweeps is further evidence of fitness costs, with no evidence of fixation in any of the populations containing sweeping haplotypes.

Although this study covers the geography of the Indo-Pacific region extensively, our investigation of *Ae. aegypti* had only single samples from each of Africa, North America, and South America. This limited our capacity to infer admixture sources and to investigate the geographical coverage of the increased copy number in New Mexico. Furthermore, while our ddRAD data provided strong evidence of sweeps and copy number variation, the sparseness of ddRAD data prevented us from inferring specific GST genes that had undergone duplication. This could be further investigated with whole genome data or specific sequencing of this region. Our analysis of *Ae. albopictus* was also limited by a lack of samples from outside the Indo-Pacific region, as well as the incompleteness of this species’ genome assembly relative to that of *Ae. aegypti*. Additional samples *Ae. albopictus* would potentially allow for similar analyses of the V1016G mutation which has been recorded recently in this species ^23^ but was not detected in any of our samples.

In summary, our analysis of *Ae. aegypti* and *Ae. albopictus* has uncovered a set of evolutionarily independent, globally segregating, partial selective sweeps at multiple resistance genes that have asynchronously spread resistance alleles across geographically distant populations. The complex patterns uncovered in these invasive disease vectors provide a detailed picture of insecticide-based selection worldwide that can help guide chemical control options for local authorities ^2,32^. These results highlight the GST epsilon class genes as having a major global impact. Our findings also indicate a set of specific resistance backgrounds to be considered in novel mosquito control strategies that rely on matched resistance levels between lab and field populations for long term success ^67^.

## Methods

### 4.1 Study Design

This study used genome-wide sequence data and endpoint genotyping assays to identify genetic backgrounds at insecticide resistance genes that showed signs of positive selection and were distributed among genomically distinct populations of *Aedes* mosquitoes . We used genetic data from 26 countries. This includes 32 populations of *Ae. aegypti* from 19 countries (n = 444) and 40 populations of *Ae. albopictus* from 17 countries (n = 490). Samples specifically sequenced for this project included *Ae. aegypti* (n = 148) from 12 populations (Kenya, Tonga, Timor-Leste, USA (New Mexico), Australia (Cape York and Torres Strait Islands) and PNG (East New Britain, East Sepik, Lae, Madang, Milne Bay, and Port Moresby)), and *Ae. albopictus* (n = 210) from 12 populations (Indonesia (Yogyakarta) and PNG (Alotau Town, Duke of York Island, East New Britain, Kokopo, Lae, Lihir Island, Malba, Mawi, North Fly Region, Vanimo, Wambisa, Wewak). South Fly Region in PNG could not be sampled due to COVID-19 restrictions on fieldwork. The *Ae. aegypti* sample from New Mexico was a lab strain in its fourth generation ^68^. All samples, including those used from previous projects, are listed in Tables S1 and S2. Genomic DNA was extracted from the 148 *Ae. aegypti* and 210 *Ae. albopictus* using Qiagen DNeasy Blood & Tissue Kits (Qiagen, Hilden, Germany) or Roche High Pure™ PCR Template Preparation Kits (Roche Molecular Systems, Inc., Pleasanton, CA, USA).

### 4.2 qPCR assays for VSSC mutations

*Aedes aegypti* were screened for F1534C, V1016G, and S989P mutations using endpoint genotyping assays described previously ^9^. For *Ae*. *albopictus*, samples were prepared for Sanger sequencing of domain III of the VSSC gene to characterise DNA sequences at codon 1534 using primers aegSCF7 (GAGAACTCGCCGATGAACTT) and aegSCR7 (GACGACGAAATCGAACAGGT) ^69^ to generate an amplicon of 740 bp. A PCR master mix was employed with final concentrations of Standard ThermoPol buffer Mg-free (1x) (New England Biolabs, Ipswich MA, USA), dNTPs (0.2 mM each) (Bioline, London UK), MgCl2 (1.5 mM) (Bioline, London UK), 0.5 µM each of forward and reverse primers, 0.625 units of Immolase™ Taq polymerase (Bioline, London, UK), 2 µL genomic DNA and PCR-grade H2O, to a final volume of 25 µL. PCR cycling conditions were: initial denaturation of 95°C for 10 min, 35 cycles of 95°C for 30s, annealing at 52°C for 45 s and extension at 72°C for 45 s, followed by a final extension of 5 min at 72°C. Amplicons were purified and sequenced by Macrogen Inc. in Seoul, Korea, on a 3730xl DNA analyser using sequencing primers Alb171F (CCGATTCGCGAGACCAACAT) ^70^ and aegSCR8 (TAGCTTTCAGCGGCTTCTTC) ^69^. Sequences were analysed using Geneious® 11.1.4 (Biomatters Ltd).

### 4.3 Sequencing and processing genomic data

Extracted DNA was used to build double digest restriction-site associated DNA (ddRAD ^71^) sequencing libraries following the protocol of Rašić, Filipović, Weeks, & Hoffmann ^72^. Libraries were individually barcoded and sequenced on either a HiSeq 4000 or a Novaseq 6000 using 150 bp chemistry. New and old sequence data were combined for each species and run through the same bioinformatic pipeline. We used the Stacks v2.54 ^73^ program “process_radtags” to demultiplex sequences and remove sequences with Phred scores below 20. Sequences were aligned to the nuclear genome assembly for *Ae. aegypti*, AaegL5 ^74^, and the linkage-based assembly for *Ae. albopictus*, AalbF3 ^75^, using Bowtie2 v2.3.4.3 ^76^ with “–very-sensitive” settings. The output .bam files were used for genomic data analysis using either Stacks (sections 4.4.1 - 4.4.3) or GATK v4.2.6.1 ^77^ (section 4.4.4).

### 4.4 Genomic data analysis

#### 4.4.1 Genetic structure and admixture

Sequences were built into Stacks catalogs for each species using the Stacks program “ref_map”. The Stacks program “populations” was used to export VCF files containing SNP genotypes for all individuals in each catalog, filtering to retain SNPs called in at least 50% of individuals from each population and 90% of individuals total, and with a minor allele count ≥ 3 ^78^. To avoid issues from low sample number, for the above filtering steps we treated PNG *Ae. aegypti* samples from East Sepik and Lae as one population, and Torres Strait Islands *Ae. albopictus* as one population (Tables S1, S2). All samples had < 30% missing data and total missing data was low (*Ae. aegypti*, x̂ = 2.42%; *Ae. albopictus*, x̂ = 4.88%). For analyses of genetic structure and latent factor mixed models, missing data were imputed in windows of 6000 SNPs using Beagle v4.1 ^79^, using 10 iterations, 500 SNP overlaps, and the full dataset as reference. Final datasets contained 79,084 SNPs for *Ae. aegypti* and 96,269 SNPs for *Ae. albopictus*.

Genome-wide genetic structure among individuals was analysed using fineRADstructure ^33^. Relevant subsets of populations were further analysed with TreeMix v1.13 ^36^ to build maximum likelihood trees and run F_3_ tests ^80^. TreeMix analysed SNPs in blocks of 1000, using a bootstrap replicate, and with the root set to Kenya (*Ae. aegypti*) or China (*Ae. albopictus*). Principal components analysis was run in R package SNPRelate v1.34.1 ^81^, using functions “snpgdsPCA” to build a principal component axis from each pair of source populations, “snpgdsPCASNPLoading” to calculate SNP loadings, and “snpgdsPCASampLoading” to project individuals onto the principal component axis.

#### 4.4.2 Genome-wide association with VSSC mutations

Latent factor mixed models associating SNP variation with VSSC genotype were run in the R package LEA v3.2.0 ^82^. Analysis of *Ae. aegypti* used all samples with available genomic and VSSC data (n = 395). Analysis of *Ae. albopictus* used all samples from the six populations where VSSC mutations were found (n = 114); analysis of all samples (n = 490) found no associations (Fig S12), potentially due to the much lower frequency of VSSC mutations in *Ae. albopictus*.

As both species exhibited strong genome-wide genetic structure among populations, we first used sparse non-negative matrix factorisation (function “snmf”) to determine an appropriate number of K clusters with which to condition the mixed models. Sparse non-negative matrix factorisation used 10 repetitions for each potential K, which identified K =18 for *Ae. aegypti* (Fig S1) and K = 4 for *Ae. albopictus* (Fig S11). Latent factor mixed models (function “lfmm”) were run using these K and a minor allele frequency of 0.05, and with 10,000 iterations, a burnin of 5000, and 10 repetitions.

Adjusted P-values were computed from the combined z-scores. Strongly associated SNPs were those that had adjusted P-values below the inverse of the number of SNPs (1.956 × 10^-5^ for *Ae. aegypti*; 2.183 × 10^-5^ for *Ae. albopictus*). Strongly-associated SNPs were found in clusters around the VSSC and GST regions; these were used in PCAs (Fig 4b, 10c) and heatmaps (Fig 6b). Principal components analyses were run in LEA (function “pca”).

#### 4.4.3 Genome-wide analysis of introgressed backgrounds

Tajima’s D, nucleotide diversity, linkage disequilibrium, and pairwise F_ST_ were calculated in VCFtools v0.1.16 using the filtered dataset without the imputation step. Tajima’s D was calculated in bins of 5 Kbp using --TajimaD 5000”. Nucleotide diversity was calculated using “--site-pi”. Linkage disequilibrium was calculated as the squared correlation coefficient between genotypes (r^2^) using “--geno-r2”, setting “--ld-window-bp-min 500” and “--ld-window-bp 100000” to ignore comparisons within RADtags or separated by more than 100 Kbp, and this was visualised using the average chromosome position and r^2^ scores of each bin. F_ST_ was calculated using “--weir-fst-pop”, “--fst-window-size 5000” and “--fst-window-step 1000”. Parameters were visualised as moving averages using the R package “tidyquant” (function “geom_ma”; ma_fun = SMA).

Linkage network analysis used VCFtools to calculate intrachromosomal (“--geno-r2”) and interchromosomal (“--interchrom-geno-r2”) linkage disequilibrium for the six subsets of individuals associated with each sweep, plus the three subsets of wild-type individuals from the same populations as the GST sweeps, plus the total set of VSSC wild types. Results were visualised with the R package “tidyverse” (function “geom_histogram”; binwidth=1). The r^2^ > 0.6 cutoff was chosen following iterative evaluation of cutoffs from 0.3 to 0.8, with r^2^ > 0.6 providing the strongest visual signal-to-noise ratio.

#### 4.4.4 Copy number variation at glutathione S-transferase (GST) genes

Copy number variation was assessed by comparing read depths at GST genes relative to those upstream and downstream. This required a genotyping and filtering pipeline that retained monomorphic as well as polymorphic sites. First, .bam files were processed in samtools v1.16 to remove unmapped reads and non-primary alignments. GVCF files were produced for each individual using HaplotypeCaller in GATK v4.2.6.1^77^, and these were genotyped using GenotypeGVCFs set to ‘--include-non-variant-sites’. Hard filtering excluded indels then followed standard GATK guidelines for non-model taxa, setting “QD < 2.0”, “QUAL < 30.0”, “SOR > 3.0”, “FS > 60.0”, and “MQ < 40.0”. Individuals were then filtered individually in bcftools v.1.16 ^83^ to remove missing data sites, sites with star alleles, and sites with less than 1X read depth. Read depths were calculated in vcftools (function “--depth”), using all sites retained after filtering. Depths at GST genes were calculated as the average depth across the 15 GST coding regions. Individuals with fewer than 500 genotyped sites within the 15 coding regions were omitted. Copy number variation was calculated at the *Ae. aegypti* VSSC gene using the same methods as above, with read depth ratios calculated from the VSSC coding region.

### 4.5 Geographical mapping

All maps were produced using ArcGIS Pro v3.1 (https://www.esri.com/en-us/arcgis/products/arcgis-pro/overview). Maps used a Mollweide projection with a central meridian of 160° E.

## Data Availability

Raw .fq files and relevant metadata for 934 mosquitoes will be available from the NCBI SRA following publication. Code used in processing, analysis, and plotting will be available from a Dryad repository. Source data will be provided with this paper, accessible via the Dryad repository.

