## Supplemental Figures S1-S12 for "Global, asynchronous partial sweeps at multiple insecticide resistance genes in *Aedes* mosquitoes"

Thomas L Schmidt *et al.*

Contents: Figs. S1 to S12

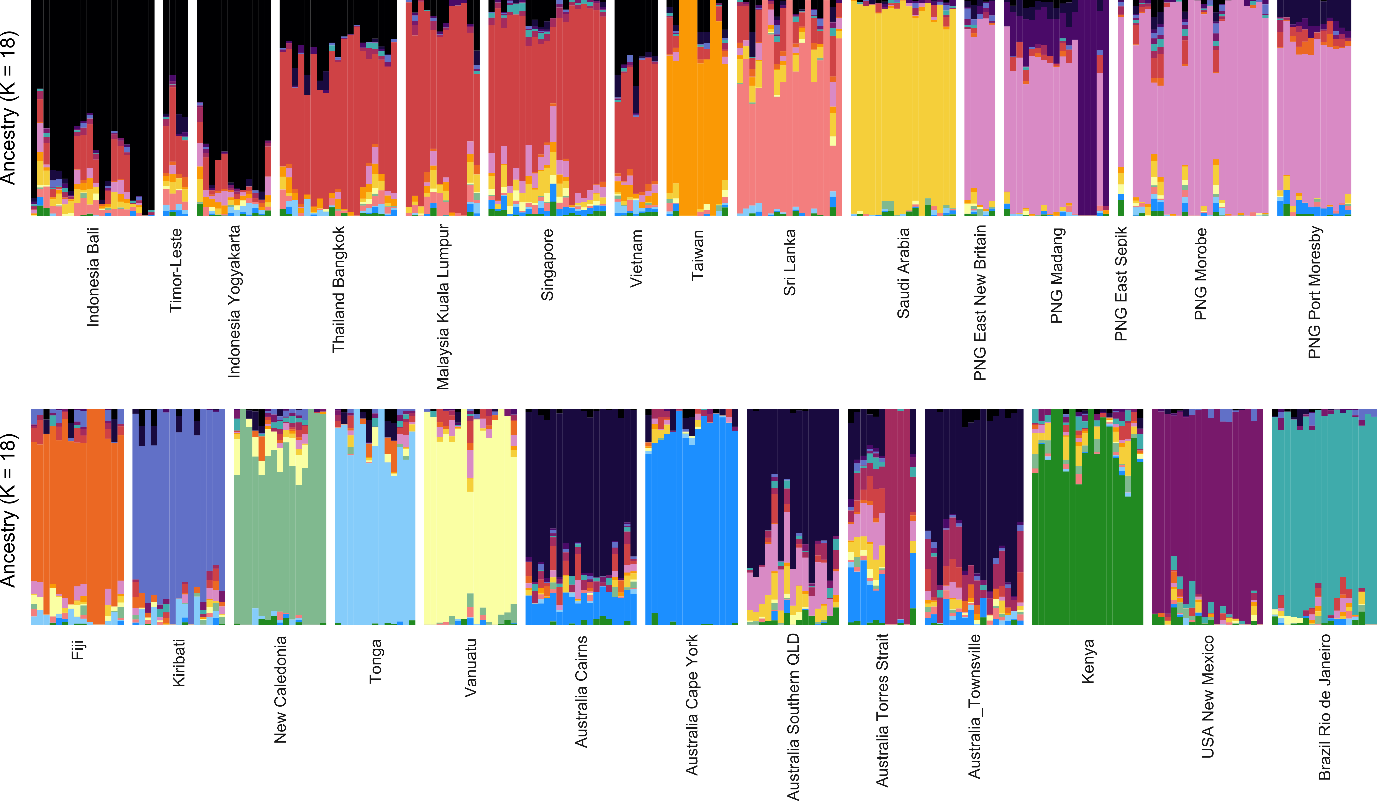

Fig S1. Sparse non-negative matrix factorisation results for *Ae. aegypti*, with K = 18. These patterns of genetic structure were used in latent factor mixed models (Figs 3b, c).

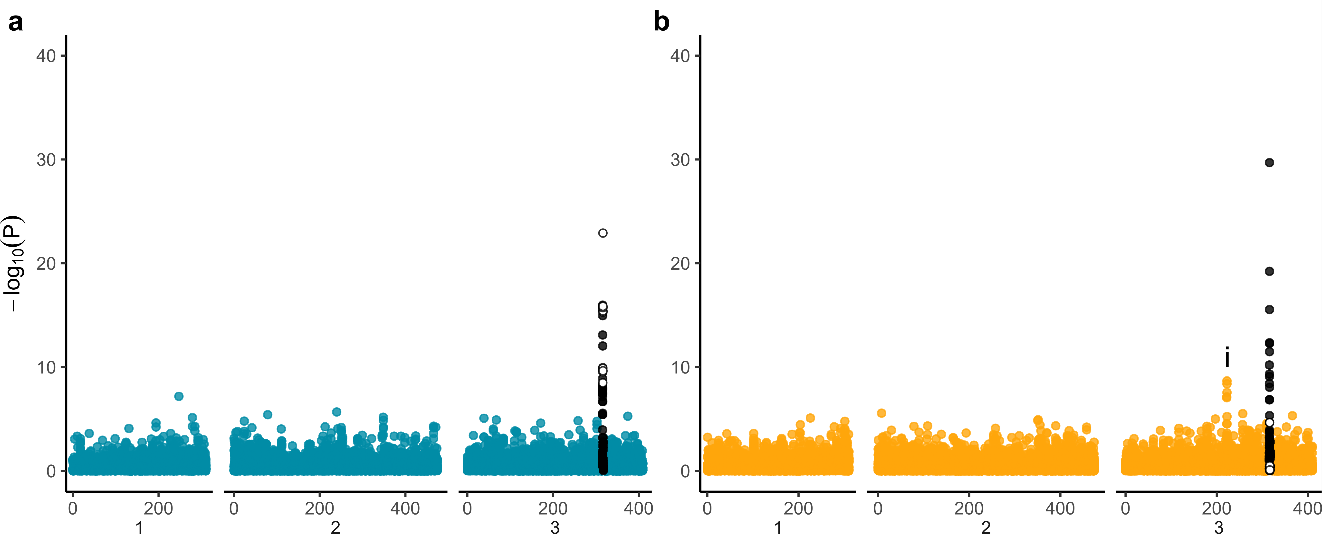

Fig S2. Latent factor mixed models associating the number of copies of the mutation under analysis with genome-wide SNPs, omitting populations with no copies of the mutation. (a) V1016G mutation, K = 9, n = 159; (b) F1534C mutation, K = 11, n = 210. White circles indicate SNPs within the VSSC gene on chromosome 3, black circles indicate SNPs within 1 Mb of this region. The ‘i’ indicates the location of a ‘Nach’ sodium channel protein (LOC5578185).

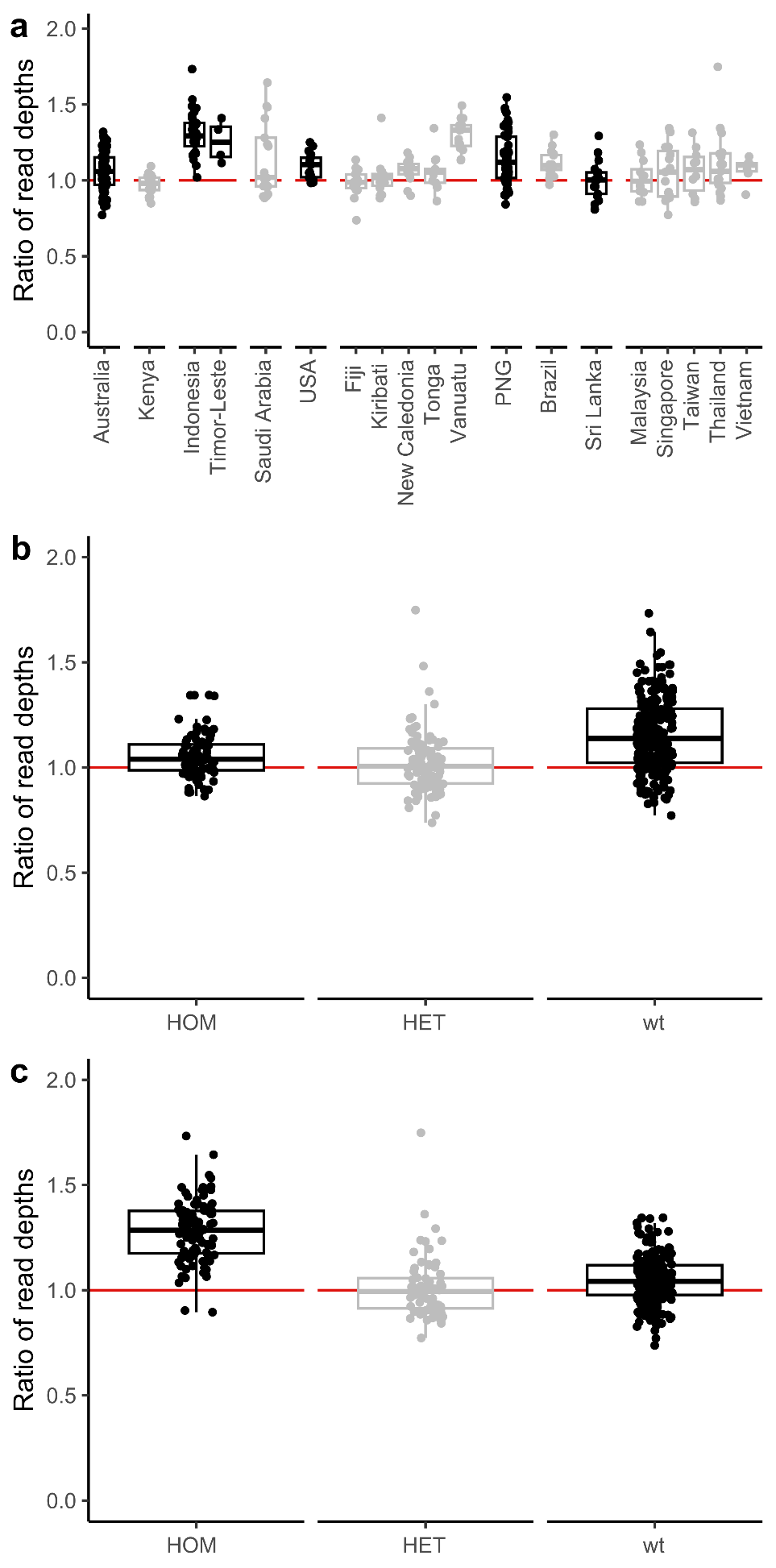

Fig S3. Ratio of read depths at the VSSC gene compared with sites <10 Mb upstream and downstream. Individuals are grouped by (a) global region and country, (b) F1534C genotype, and (c) V1016G genotype.

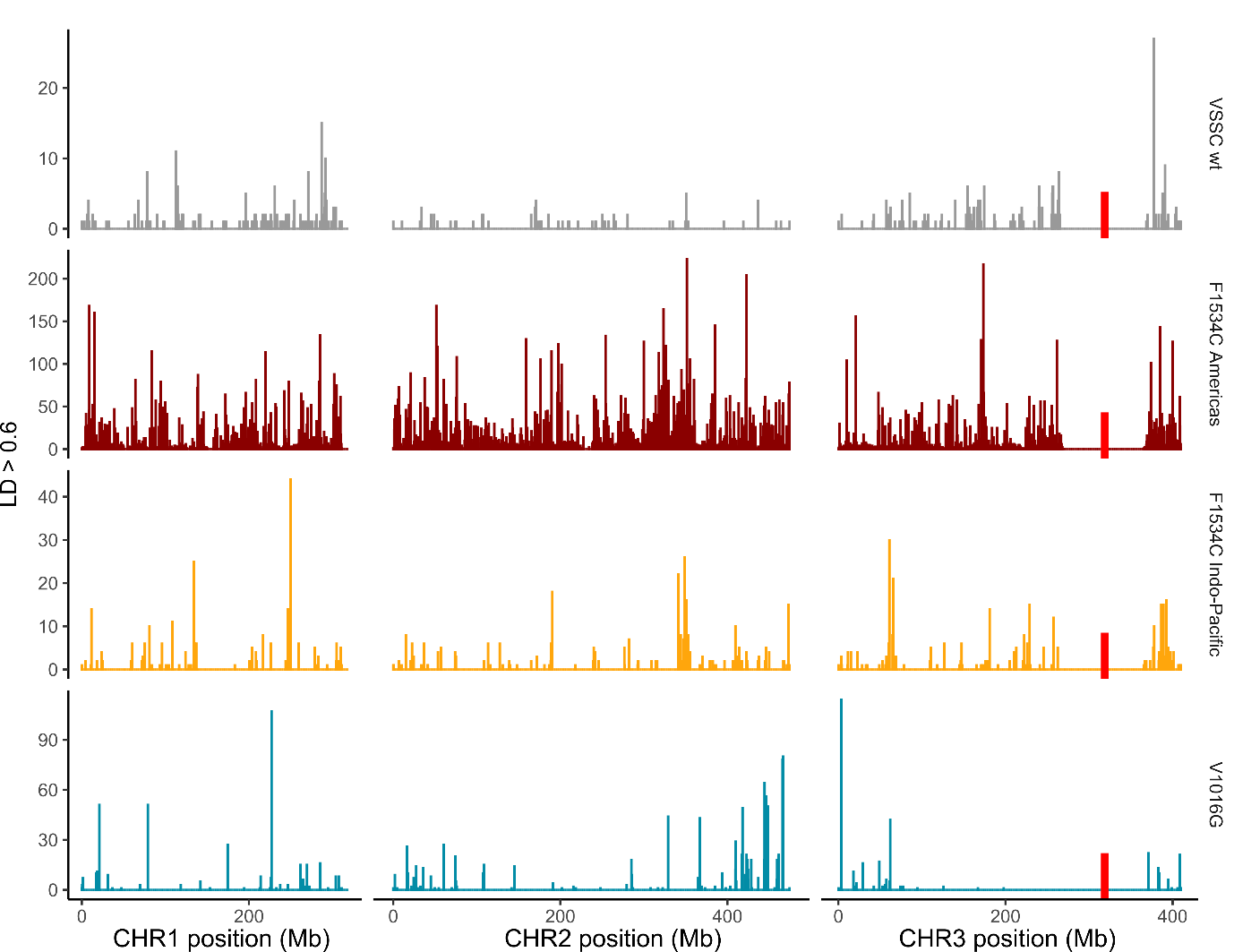

Fig S4. Linkage network analysis in *Ae. aegypti* VSSC wild types. Rows indicate the VSSC wild-type individuals (grey) and the three VSSC backgrounds (colours). Plots are histograms with 500 Kb bins, showing locations of SNPs with r^2^ > 0.6 to at least one SNP within 1 Mb of the sweep locus (red bars), and scoring SNPs for each r^2^ > 0.6 interaction with a SNP near the locus. SNPs within 50 Mb of the sweep locus were omitted.

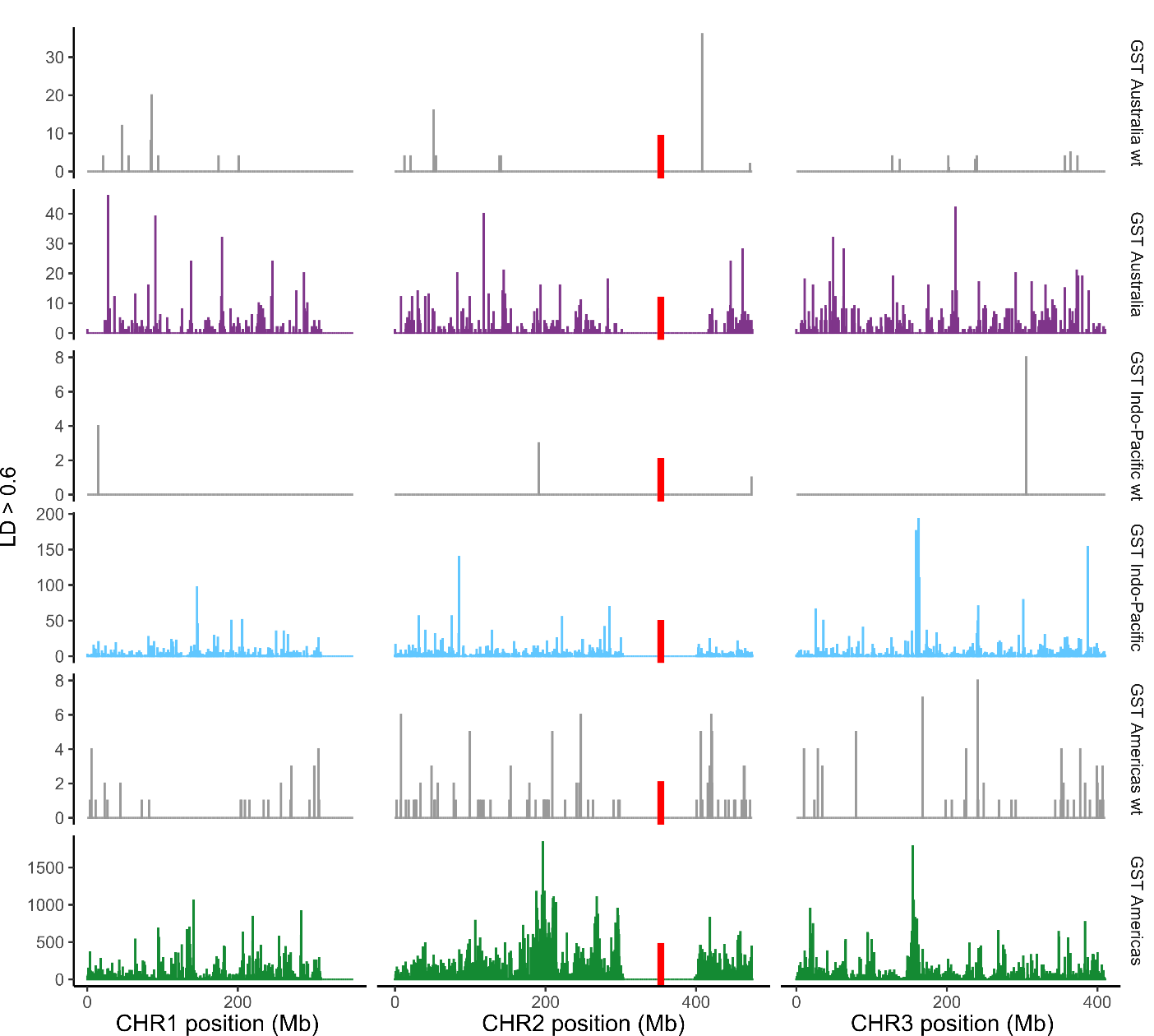

Fig S5. Linkage network analysis in *Ae. aegypti* GST wild types. Rows indicate the GST wild-type individuals (grey) and the three VSSC backgrounds (colours). Plots are histograms with 500 Kb bins, showing locations of SNPs with r^2^ > 0.6 to at least one SNP within 1 Mb of the sweep locus (red bars), and scoring SNPs for each r^2^ > 0.6 interaction with a SNP near the locus. SNPs within 50 Mb of the sweep locus were omitted.

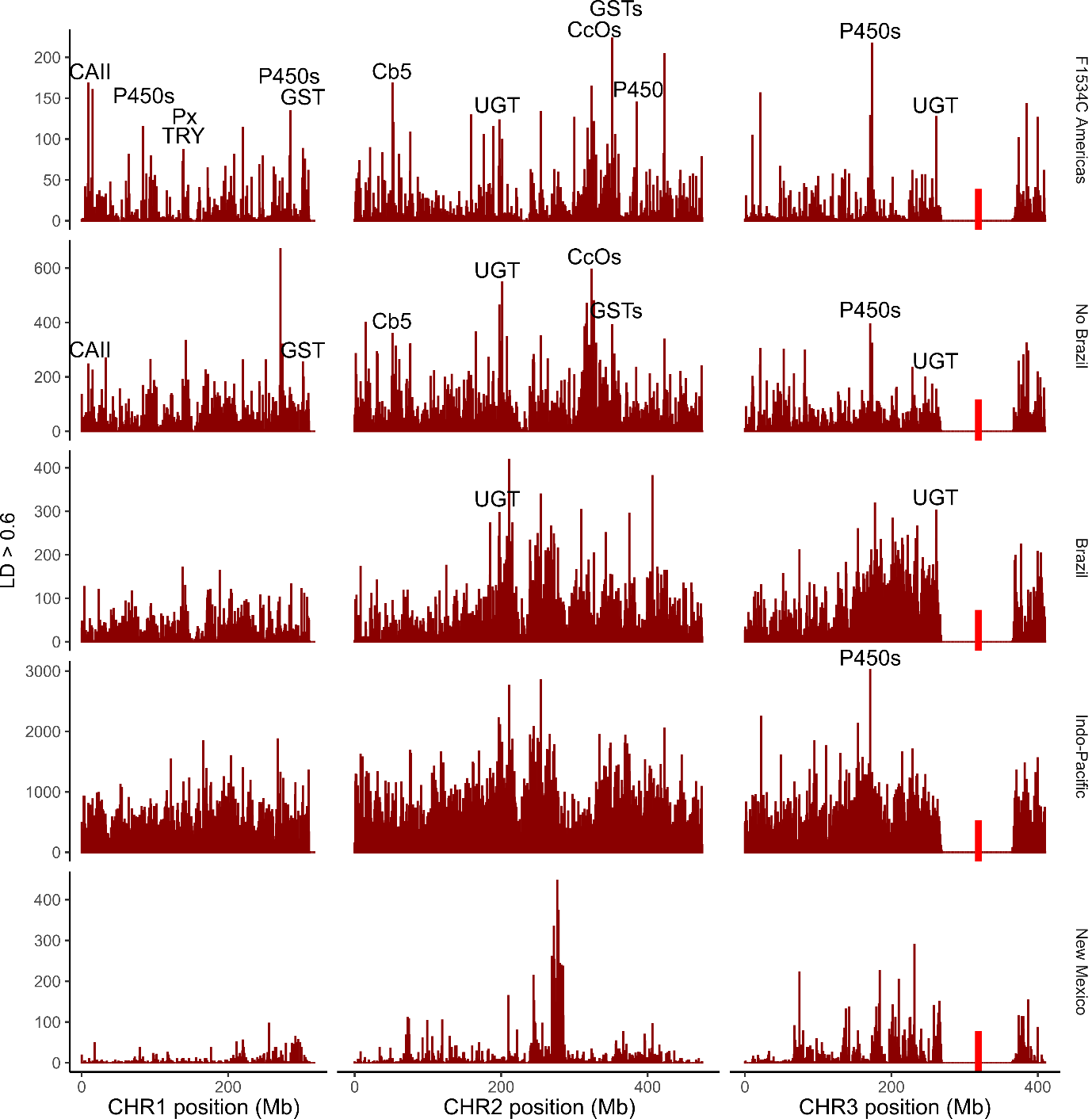

Fig S6. Linkage network analysis in *Ae. aegypti* F1534C Americas background. Rows indicate the F1534C Americas background (top) and subsets of individuals with the background. Plots are histograms with 500 Kb bins, showing locations of SNPs with r^2^ > 0.6 to at least one SNP within 1 Mb of the sweep locus (red bars), and scoring SNPs for each r^2^ > 0.6 interaction with a SNP near the locus. SNPs within 50 Mb of the sweep locus were omitted. Text follows the same key as Fig 9, and on subset rows text indicates peaks common to both the full dataset and the subset.

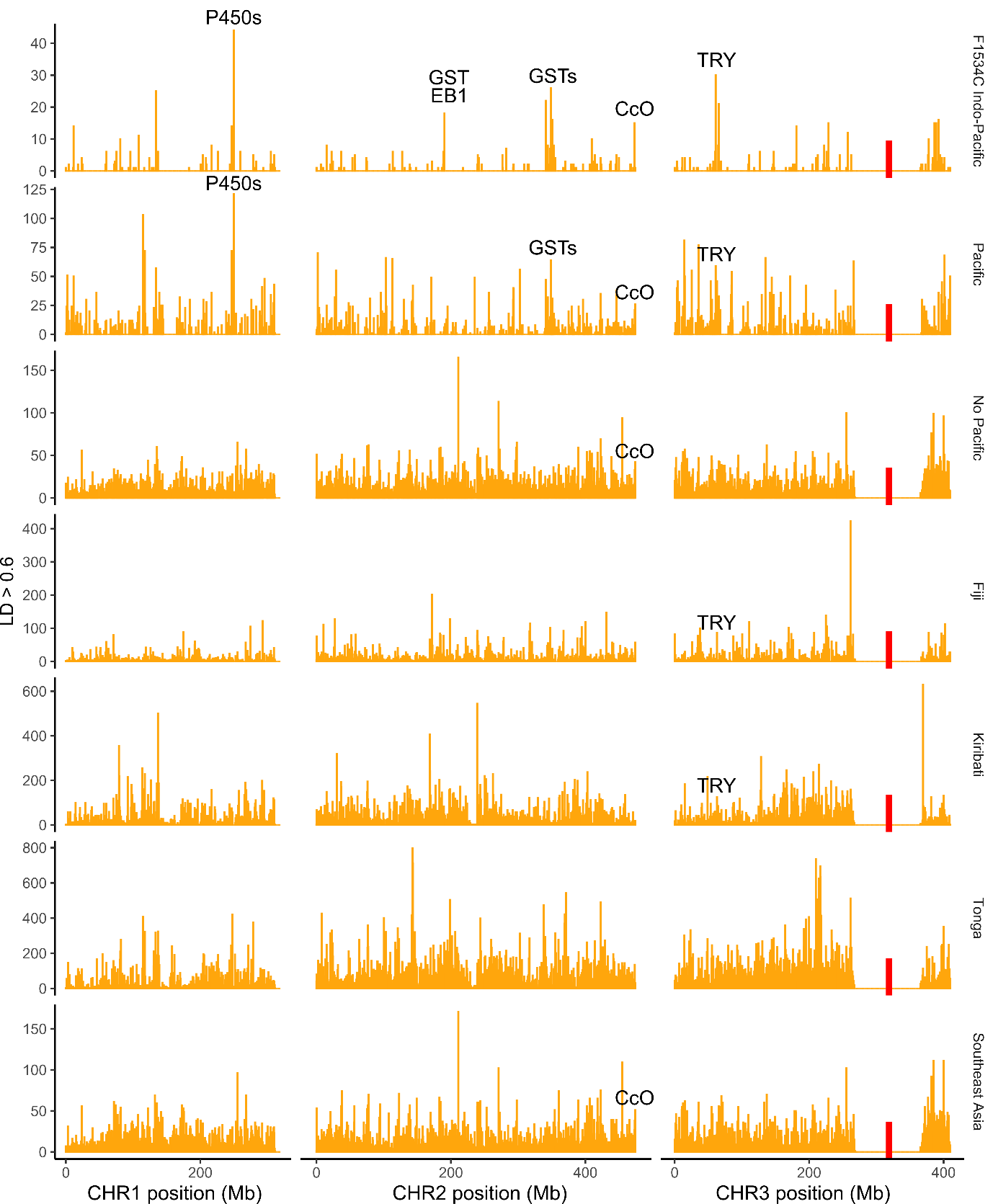

Fig S7. Linkage network analysis in *Ae. aegypti* F1534C Indo-Pacific background. Rows indicate the F1534C Indo-Pacific background (top) and subsets of individuals with the background. Plots are histograms with 500 Kb bins, showing locations of SNPs with r^2^ > 0.6 to at least one SNP within 1 Mb of the sweep locus (red bars), and scoring SNPs for each r^2^ > 0.6 interaction with a SNP near the locus. SNPs within 50 Mb of the sweep locus were omitted. Text follows the same key as Fig 9, and on subset rows text indicates peaks common to both the full dataset and the subset.

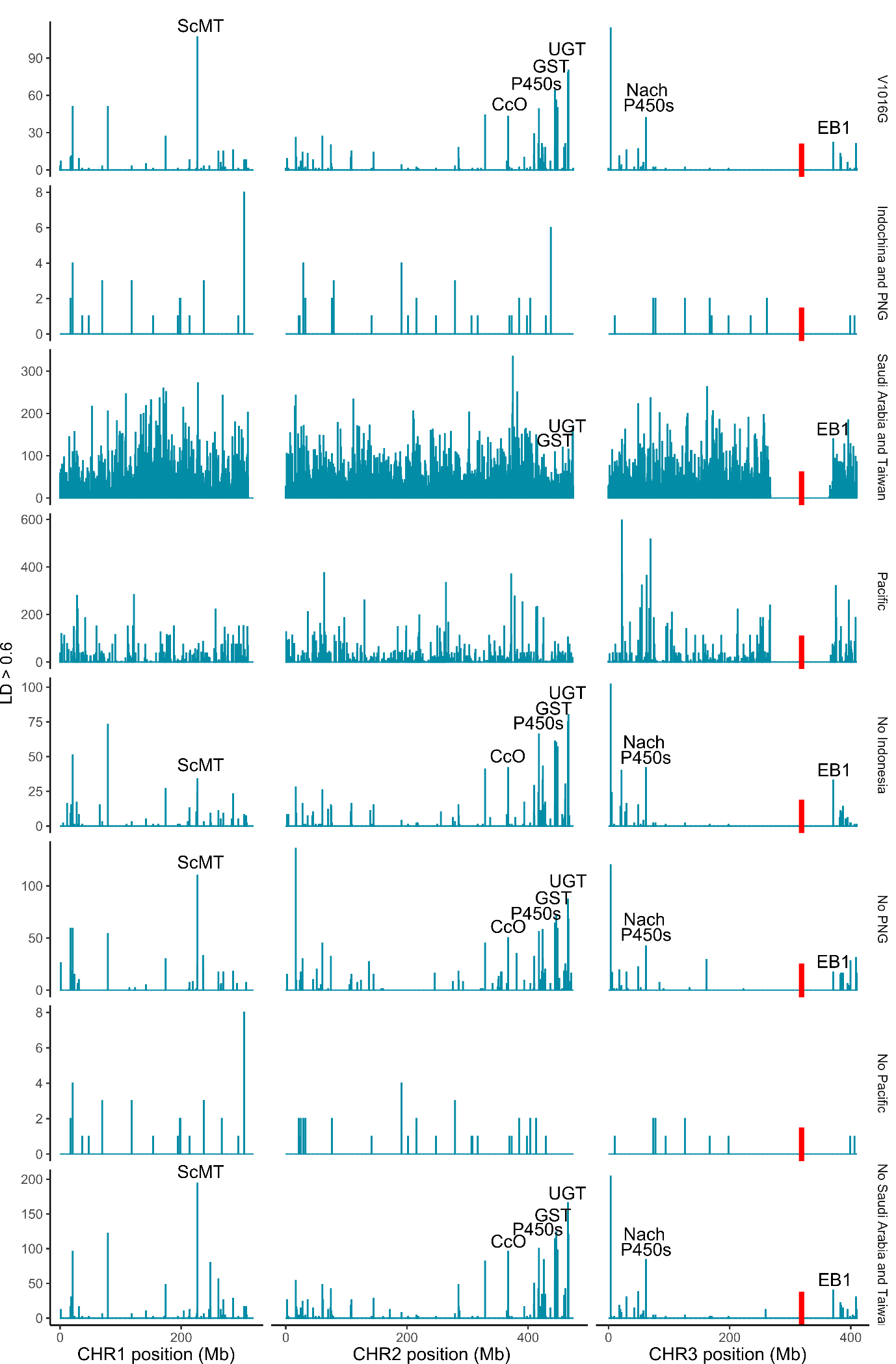

Fig S8. Linkage network analysis in *Ae. aegypti* V1016G background. Rows indicate the V1016G background (top) and subsets of individuals with the background. Plots are histograms with 500 Kb bins, showing locations of SNPs with r^2^ > 0.6 to at least one SNP within 1 Mb of the sweep locus (red bars), and scoring SNPs for each r^2^ > 0.6 interaction with a SNP near the locus. SNPs within 50 Mb of the sweep locus were omitted. Text follows the same key as Fig 9, and on subset rows text indicates peaks common to both the full dataset and the subset.

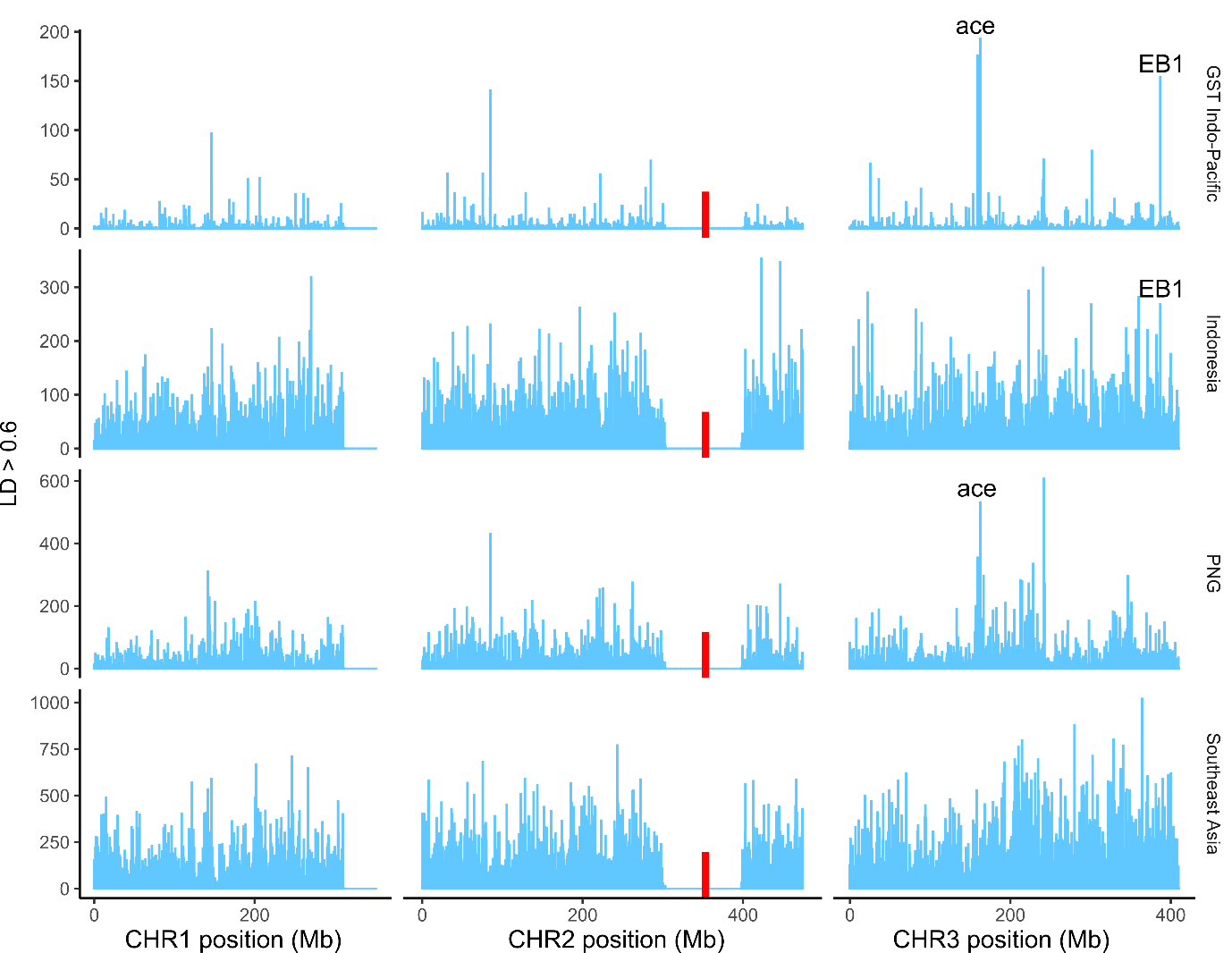

Fig S9. Linkage network analysis in *Ae. aegypti* GST Indo-Pacific background. Rows indicate the GST Indo-Pacific background (top) and subsets of individuals with the background. Plots are histograms with 500 Kb bins, showing locations of SNPs with r^2^ > 0.6 to at least one SNP within 1 Mb of the sweep locus (red bars), and scoring SNPs for each r^2^ > 0.6 interaction with a SNP near the locus. SNPs within 50 Mb of the sweep locus were omitted. Text follows the same key as Fig 9, and on subset rows text indicates peaks common to both the full dataset and the subset.

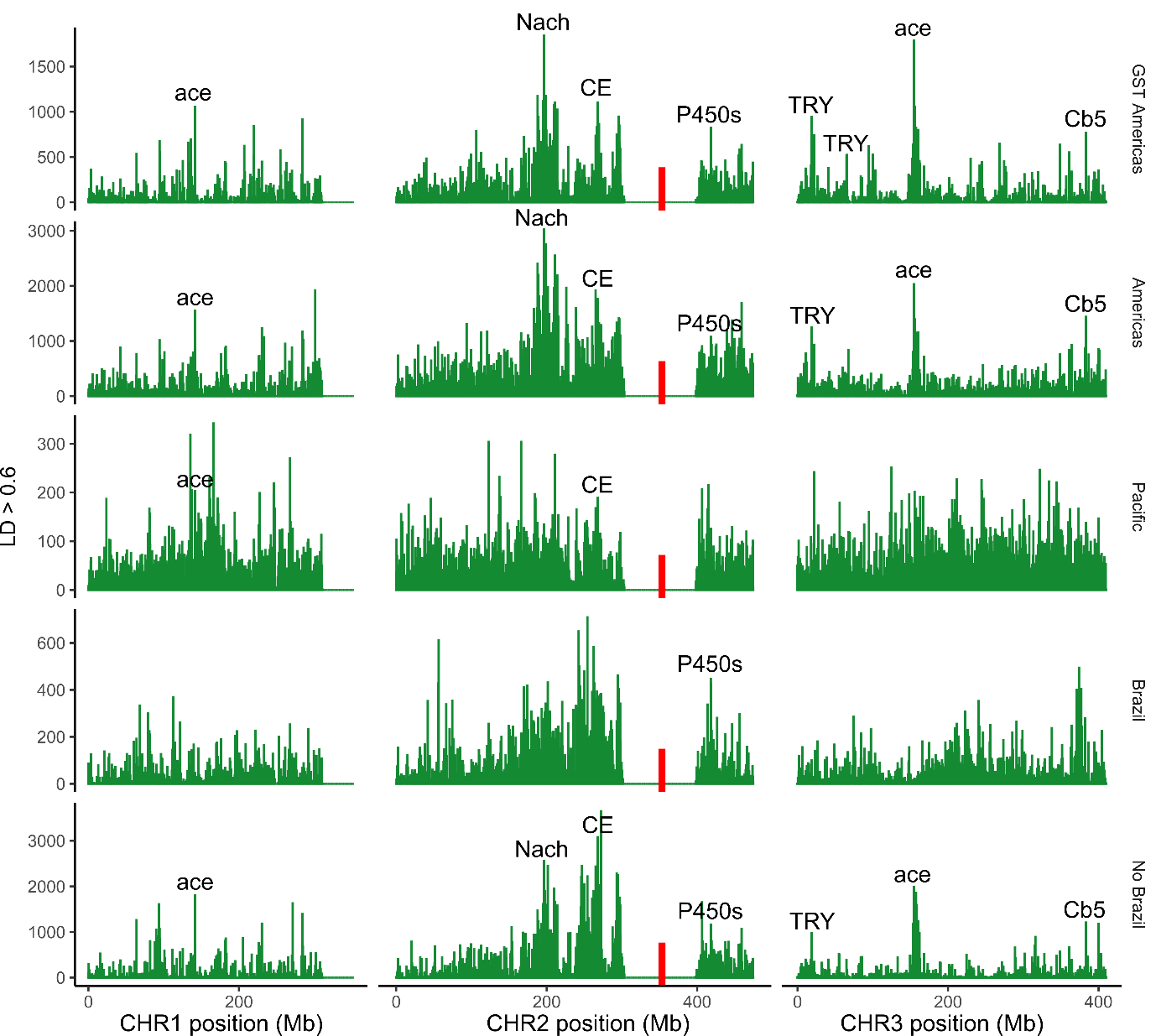

Fig S10. Linkage network analysis in *Ae. aegypti* GST Americas background. Rows indicate the GST Americas background (top) and subsets of individuals with the background. Plots are histograms with 500 Kb bins, showing locations of SNPs with r^2^ > 0.6 to at least one SNP within 1 Mb of the sweep locus (red bars), and scoring SNPs for each r^2^ > 0.6 interaction with a SNP near the locus. SNPs within 50 Mb of the sweep locus were omitted. Text follows the same key as Fig 9, and on subset rows text indicates peaks common to both the full dataset and the subset.

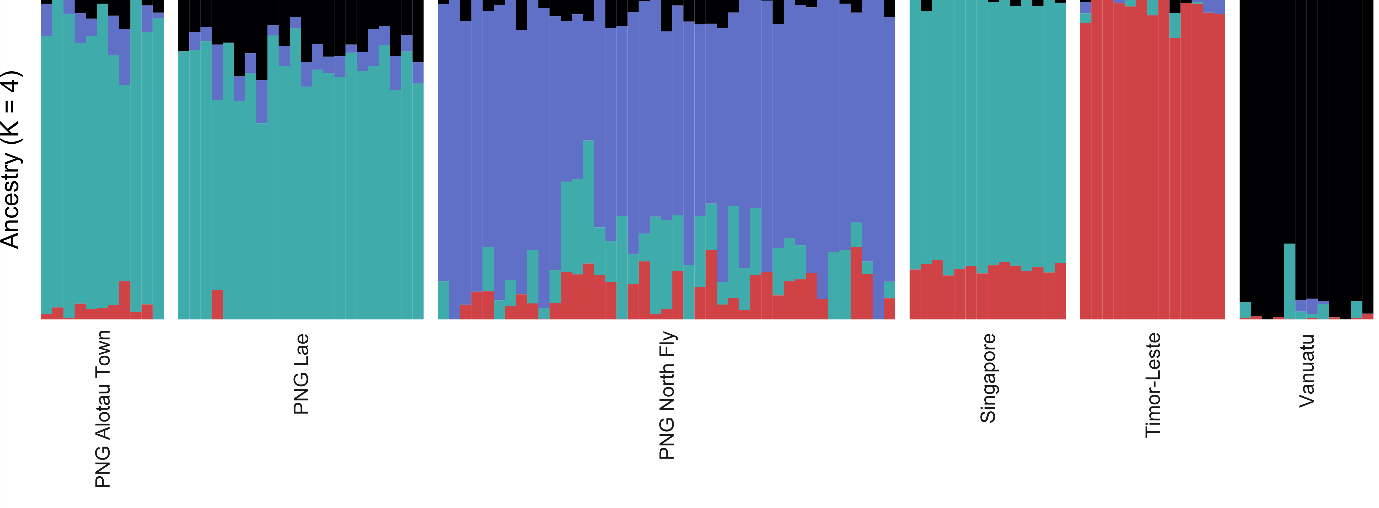

Fig S11. Sparse non-negative matrix factorisation results for *Ae. albopictus*, setting K = 4. These patterns of genetic structure were used in the latent factor mixed model (Fig 10b).

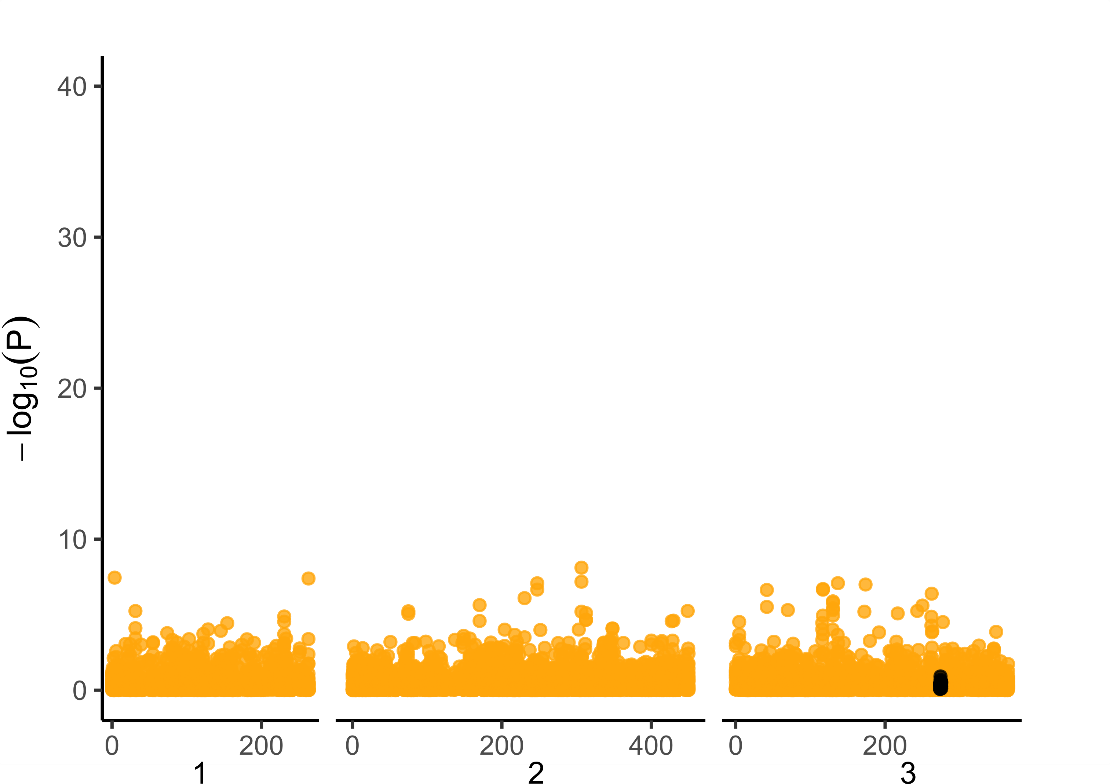

Fig S12. Latent factor mixed model associating the number of copies of the F1534C mutation with genome-wide SNPs, using *Ae. albopictus* from all populations (n = 490). Black circles indicate SNPs within 1 Mb of the VSSC gene on chromosome 3. Model is conditioned on K = 10.
